# Intron retention regulates STAT2 function and predicts immunotherapy response in lung cancer

**DOI:** 10.1101/2025.08.19.671121

**Authors:** Ryan P. Englander, Mattia Brugiolo, Te-Chia Wu, Mitch Kostich, Nathan K. Leclair, SungHee Park, Jacques Banchereau, Peter Yu, Andrew Salner, Romain Banchereau, Karolina Palucka, Olga Anczuków

## Abstract

Immunotherapy benefits only a subset of lung cancer patients, and the molecular determinants of variable outcomes remain unclear. Using long-read RNA-sequencing we mapped the landscape of alternative RNA splicing in human primary lung adenocarcinomas. We identified over 180,000 full-length mRNA isoforms, more than half of which were novel and many of which occurred in immune-related genes, particularly within the type I interferon response pathway. We discovered retained introns in the *STAT2* gene that produce altered protein isoforms that regulate immune signaling and interferon responses. Furthermore, *STAT2* intron retention levels predicted patient responses to checkpoint inhibitors. These findings position alternative splicing as a key regulator of tumor immune responses and reveal mRNA splicing alterations as potential biomarkers to identify patients who might respond to cancer immunotherapy.

## Introduction

Lung cancer is a leading cause of death globally, responsible for almost 2 million deaths in 2018 (*1*). Lung adenocarcinoma (LUAD), the most prevalent subtype, accounts for 40% of new cases (*2*). While immune checkpoint inhibitors have improved disease prognosis, many patients do not benefit, and existing biomarkers fail to predict responses adequately (*3–5*). Better understanding the molecular pathways regulating immune function in LUAD is vital for improving immunotherapy.

Alternative RNA splicing, a ubiquitous post-transcriptional process that generates multiple mRNA from a single gene (*6*), may be key to a more precise understanding of immune responses in tumors. Although almost all multiexonic genes are alternatively spliced in humans, most isoforms remain uncharacterized (*7, 8*). Splicing is often altered in cancer to favor oncogenic and immune-evasive isoforms (*9–12*), but the mechanisms underlying these changes in LUAD are poorly understood. Long-read RNA sequencing (LR-seq) allows sequencing of full-length transcripts and thus enables deeper functional analyses than would be possible utilizing conventional short-read RNA-sequencing (SR-seq) technology (*13–18*). However, to date LR-seq has only been applied to studies of alternative splicing in LUAD cell lines and not primary tumors (*19–21*). In primary tumors from other cancer types, LR-seq has revealed thousands of novel isoforms (*22–25*), highlighting its potential to uncover biologically and clinically relevant isoforms in LUAD.

Using LR-seq and a multilevel analytical platform, we mapped full-length mRNA isoforms in LUAD tumors. Many novel isoforms impacted genes in immune pathways, notably interferon (IFN) signaling. IFNs—including type I (IFN-I), III (IFN-III), and IFN-gamma—play dual roles in antitumor immunity and immune evasion (*26–36*). We identified and characterized two novel isoforms in the *STAT2* gene, an essential transcription factor necessary for transducing IFN-I/III responses. These events impair IFN responsiveness, are tumor-enriched, are targetable by RNA-based methods, and predict patient survival after PD-L1 inhibition. Our findings underscore LR-seq’s utility in discovering novel immune-regulatory mechanisms in cancer.

### LR-seq reveals thousands of novel unannotated alternative spliced isoforms in human lung tumors

To interrogate the alternative splicing landscape of human lung cancer, we performed SR-seq and LR-seq on bulk RNA from 12 LUAD tumors (**Table S1A-C**) using PacBio single-molecule real time (SMRT) circular consensus sequencing (**Fig. 1A**). Isoforms were assembled using the IsoSeq pipeline, aligned, refined, and merged into a unified transcriptome (*37, 38*). We validated splice junctions and transcript boundaries using orthogonal datasets and categorized isoforms by structure (**See Methods**).

**Figure 1.**
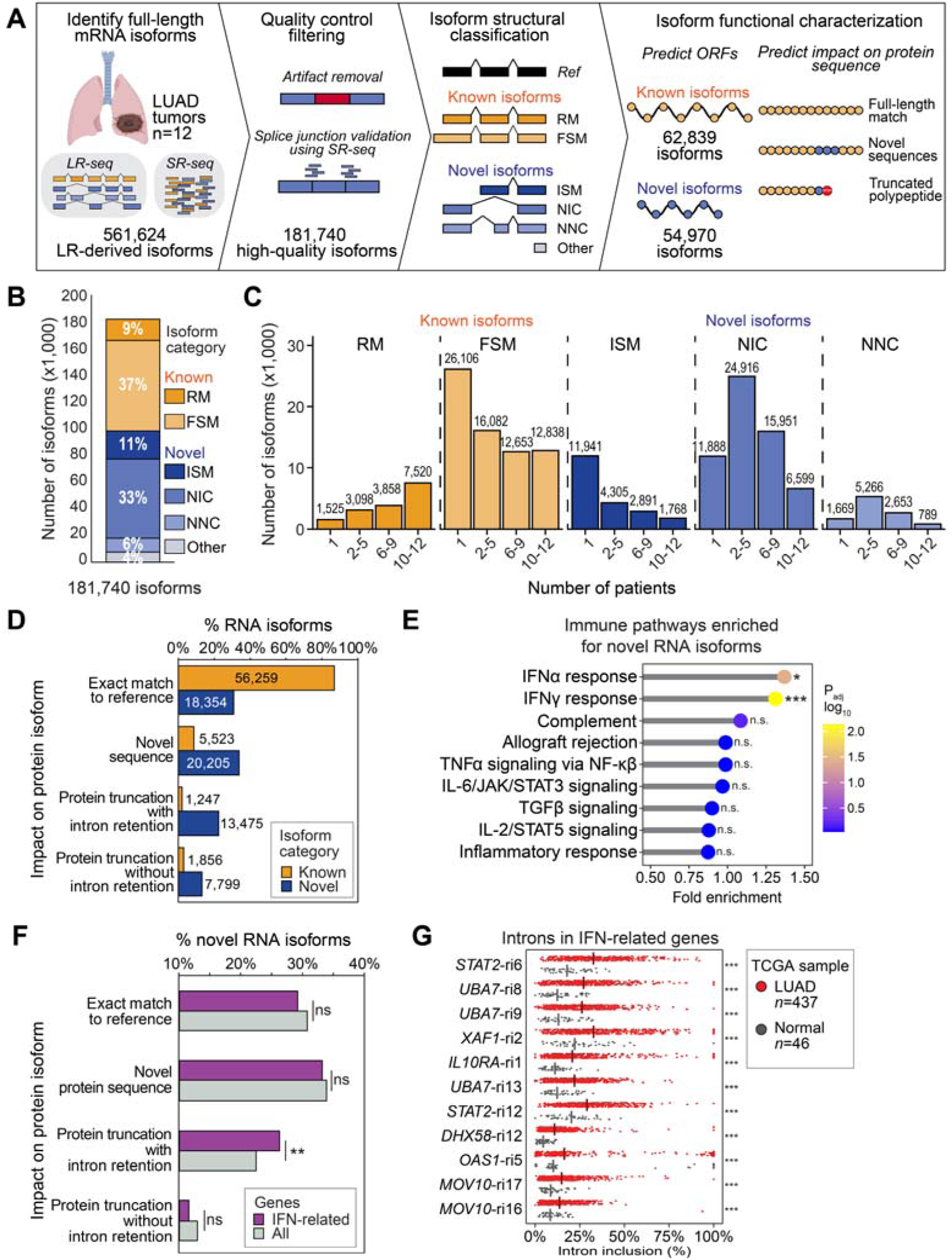
LR-seq detects novel unannotated alternatively spliced isoforms in LUAD primary tumors that are enriched in interferon-associated genes. (A) Pipeline for the discovery of full-length spliced mRNA isoforms in human primary LUAD tumors. RNA from each tumor is sequenced with LR-seq to identify novel isoforms, which undergo extensive quality controlling and filtering including SR-seq data from the same sample to validate novel spliced junctions. Full-length isoforms are annotated using SQANTI3 isoform structural categories: FSM-full-splice match; RM-reference match; ISM-Incomplete-splice match; NIC-novel-in-catalog; NNC-novel-not-in-catalog. (B) SQANTI3 isoform structural categories of LR-derived full-length mRNA isoforms detected in LUAD. (C) Patient samples positive for each isoform as quantified by FLAIR, shown by isoform structural category. (D) The impact of LR-derived full-length mRNA isoforms on protein coding potential is predicted by comparing the *in silico* translated mRNA sequence to the reference protein sequence from UniProt and separated by ‘Known’ and ‘Novel’ isoforms. See **Methods** for more details. (E) Pathway enrichment analysis of genes with more novel than known isoforms using immune-related Hallmark gene sets from MSigDB. See **Table S2F** for genes names and isoform numbers. (F) The impact of LR-derived mRNA isoforms on the protein coding potential is predicted for novel LR-derived isoforms impacting genes associated with IFNα and IFNγ response pathways (n=1,222) vs. all novel LR-derived isoforms (n=59,833) (Hypergeometric test, \**P*<0.05, \*\**P*<0.01, \*\*\**P*<0.001, ns-not significant). (G) Intron retention events are quantified as percent spliced-in (PSI) using SR-seq data from TCGA LUAD tumor and normal lung samples. Intron retention events are filtered for those supported by LR-seq and skipped in the canonical reference isoform, ranked by the difference in median PSI between tumor and matched normal samples (FDR-corrected p-values from rMATS, \**P*<0.05, \*\**P*<0.01, \*\*\**P*<0.001, ns-not significant).

After stringent filtering, we identified 181,740 full-length mRNA isoforms from 21,070 loci (**Figs. 1B** and **S1A**), with 54% novel isoforms unannotated in the GENCODE reference transcriptome (**Fig. 1B** and **Table S2A**). Isoform quantification across tumors showed consistent structural category proportions between patients (**Fig. S1B** and **Table S2B**). Notably, 47% of isoforms (n=25,992) from the novel-in-catalog (NIC) and novel-not-in-catalog (NNC) categories were found in ≥50% of patients, suggesting these novel isoforms are recurrent components of LUAD transcriptomes rather than patient-specific (**Fig. 1C** and **Table S2B**). Furthermore, full-splice-match (FSM) isoforms that exactly match splice junctions with a reference isoform exhibited shorter 3’ UTRs, (**Fig. S1C** and **Table S2C**), consistent with 3’ UTR shortening previously described in cancer (*39*). Additionally, 2% (3,676) of isoforms arose from intergenic regions, with 36% of these being multi-exonic (**Fig. S1D**), potentially representing novel or re-expressed genes.

We then compared predicted isoform protein coding sequences for LR-derived isoforms to human reference proteins from UniProtKB/Swiss-Prot (*40*). Most reference-matching isoforms had exact protein matches (78% reference-match (RM), 72% FSM), while novel structural categories frequently encoded divergent proteins (**Fig. S1E-J** and **Table S2D**). Thus, a substantial proportion of the protein-coding transcriptome in LUAD remains uncharacterized.

To determine how novel LR-derived isoforms might contribute to the LUAD proteome, we classified isoforms from known genes and their predicted open reading frames (ORFs) as either known, novel, or truncated relative to the reference (**Table S2E**). We further classified isoforms predicted to encode truncated ORFs into those with or without intron retention events, which often lead to ORF truncation by shifting the reading frame or introducing in-frame premature termination codons (PTCs) (*41*). Strikingly, out of the 59,833 novel LR-derived isoforms that could be annotated at the protein level, 36% encoded truncated proteins, 34% encoded novel proteins, and 31% encoded a known ORF (**Fig. 1D** and **Table S2E**). These data show that unannotated isoforms that can give rise to novel and truncated proteins are prevalent in LUAD.

### Novel isoforms are enriched in interferon-related genes

Despite their ubiquity, most alternatively spliced isoforms are functionally uncharacterized (*7, 42*). We chose to focus on the role of novel LR isoforms on immune-related pathways, which are critical for tumor immunity and the target of immune-modulating therapies (*5, 43–48*). Intriguingly, pathway analysis of genes enriched in novel isoforms revealed significant enrichment in IFN-alpha and IFN-gamma responses (**Fig. 1E**). We found novel isoforms in 180 of 217 IFN-related genes, including *STAT1*, *STAT2*, and *HLA-A* (**Table S2G**), with 26% (*n*=321) of novel isoforms in IFN-related genes predicted to encode truncated ORFs due to intron retention—higher than the overall novel isoform average (**Fig. 1F**).

To further explore tumor-upregulated retained introns in IFN-related genes, we analyzed SR-seq from 437 LUAD tumors and 46 normal tissues from TCGA, using GENCODE v44 supplemented with our LR-derived isoforms and quantifying splicing with rMATS (**Fig. S1K**). We found 3,151 differential splicing events (|ΔPSI| >5%, FDR<0.05) across 1,703 genes, including 73 events in 32 IFN-related genes (**Fig. S1L,M** and **Table S2H**). Filtering for intron retention in IFN-related genes yielded 11 LR-validated events in 7 genes (**Table S2I**). These events include: i) two intron retention events in *STAT2*, the prototypical IFN-I/III transcription factor (*49–52*); ii) three intron retention events in *UBA7*, an enzyme that covalently modifies to promote IFN responses and tumor immunity (*53–55*); iii) an intron retention event in *XAF1*, a tumor suppressor that enhances IFN-related apoptosis (*56, 57*); and iv) intron retention events in *DHX58*, *OAS1*, and *MOV10*, all of which are antiviral effectors with complex roles in tumor immunity (*58–63*) (**Fig. 1G** and **Table S2I,H**). Since intron retention generally reduces protein production by introducing PTCs or preventing mRNA export, these results suggest that intron-retaining isoforms disrupt immune signaling in LUAD.

### *STAT2* isoforms with retained introns are expressed in tumor cells

We focused on unravelling the role and regulation of the retained introns 6 (ri6) and 12 (ri12) in *STAT2*, as these events encompassed the most upregulated intron retention event in TCGA and *STAT2* plays a crucial role in transducing IFN-I/III responses (*26, 50, 51, 53*). LR-seq revealed 25 *STAT2* isoforms, 18 of which were novel, and 12 of which retained ri6, ri12, or both (**Fig. 2A** and **Table S2G**). We quantified the frequency of each intron retention event across our 12 LUAD patients and TCGA-LUAD and found that retention of ri6 and ri12 was correlated in most patients (**Fig. 2B,C**).

**Figure 2.**
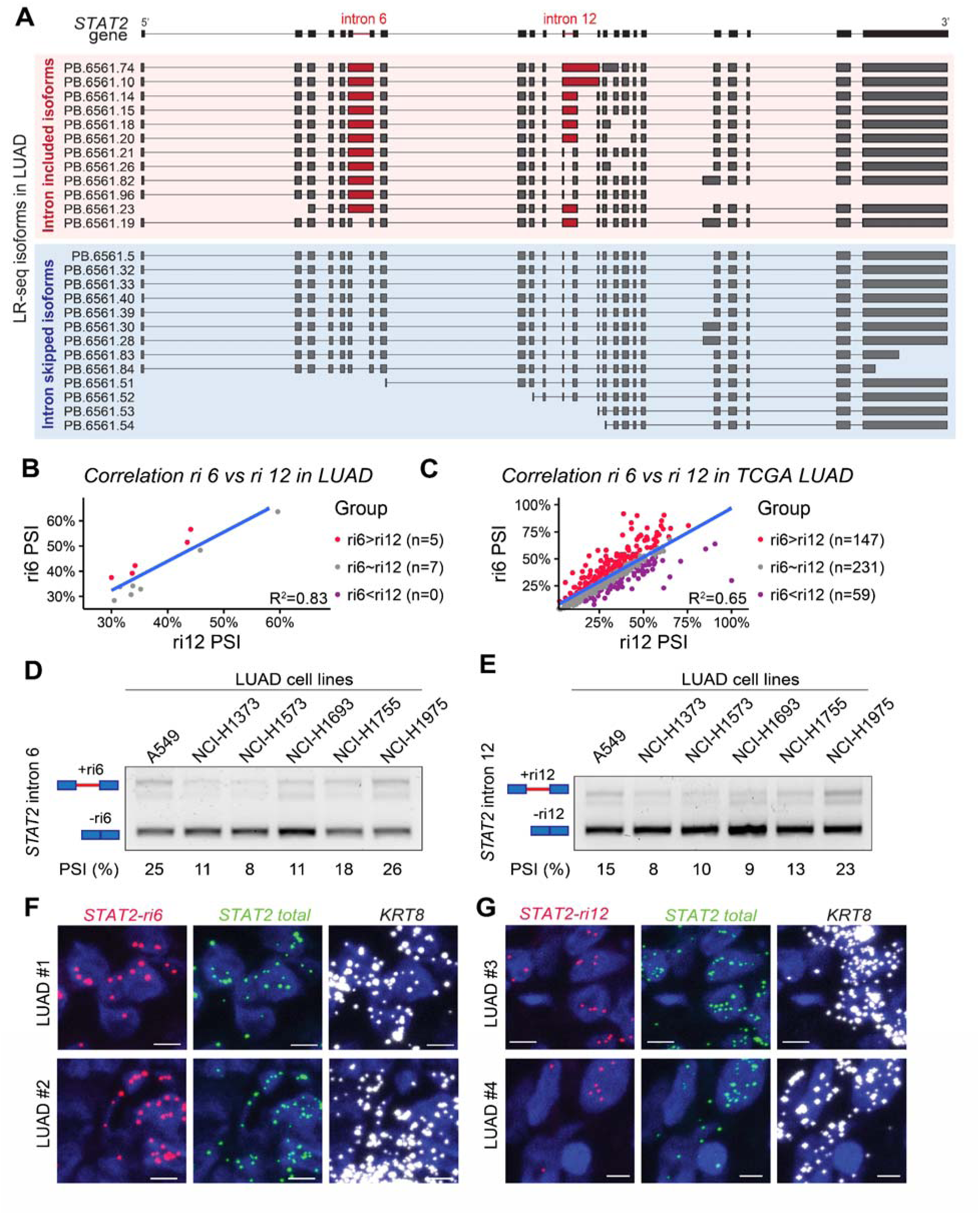
*STAT2* retained introns are expressed in cancer cell lines and primary human tumors. **(A**) Full-length spliced mRNA isoforms *STAT2* detected by LR-seq in LUAD samples, grouped by intron retained (middle) and intron skipped (bottom) isoforms. Full gene structure is shown on top with ri6 and ri12 highlighted. **(B,C)** Correlation between retention of ri6 and ri12 from SR-seq data from LUAD tumors (*n*=12) and TCGA-LUAD (*n*=437). Tumors are grouped as follows: ri6>ri12 (ri6 PSI - ri12 PSI ≥5%), ri6∼ri12 (5% > ri6 PSI - ri12 PSI >-5%), or ri6<ri12 (ri6 PSI - ri12 PSI ≤-5%). **(D,E)** Intron retention levels from ri6 (D) and ri12 (E) in six LUAD cell lines as assessed by RT-PCR with primers located in exons 6 and 9 or 8 and 13 respectively, quantified as PSI. **(F,G)** *STAT2* intron 6 (F) or intron 12 (G) containing mRNAs are visualized by RNA-FISH in LUAD tumor sections using probes targeting *STAT2*-ri6, *STAT2*-ri12 or *STAT2*-total mRNAs, as well as *KRT8* as a marker or epithelial tumor cells, with nuclei counter stained with DAPI. Scale bar 5 μm.

To confirm that *STAT2* ri6/ri12 were retained in tumor cells directly and not exclusive to other components of the tumor microenvironment, we performed RT-PCR on RNA from six LUAD cell lines using primers that amplify both the intron-containing and fully spliced mRNAs for both introns (**Fig. 2D,E**). While all cell lines predominantly express the fully spliced *STAT2* mRNAs, intron retention ranged from 8-26% for ri6, and 8-23% for ri12 (**Fig. 2D,E**). RNA fluorescence *in situ* hybridization (FISH) on LUAD tumor sections using probes targeting *STAT2* ri6, *STAT2* ri12, *STAT2* total mRNAs, and the epithelial tumor cell marker *KRT8* showed signal for *STAT2* ri6 and ri12 in *KRT8*+ tumor cells (**Fig. 2F,G**), demonstrating that *STAT2* intron retention occurs in LUAD cell lines as well as cancer cells from patient tumors.

### Antisense oligonucleotide-mediated modulation of *STAT2* intron retention impacts STAT2 protein expression and interferon signaling

To assess how *STAT2* intron retention affects protein expression and IFN signaling, we designed splice-switching antisense oligonucleotides (ASOs) targeting the 5’ or 3’ splice sites (5’/3’SS) of *STAT2* ri6 or ri12 (**Fig. 3A,B**). In A549 lung cancer cells, ASOs either decreased or increased retention of targeted introns, with largely consistent effects at baseline and after IFN-I stimulation (**Fig. 3C-H**). For ri6-targeting ASOs, 3 out of 4 decreased intron 6 retention (ASOs ri6-5’SS1, ri6-3’SS1 and ri6-3’SS2) (**Fig. 3C**), with ASOs ri6-3’SS1 and ri6-3’SS2 also decreasing intron 12 retention (**Fig. 3E**) and trending towards or significantly increasing STAT2 protein levels at baseline and upon IFN-I stimulation, respectively (**Fig. 3G**). By contrast, ASO ri6-5’SS2 increased intron 12 retention and decreased STAT2 protein expression by >10-fold (**Fig. 3C,E,G**). For ri12-targeting ASOs, ASOs ri12-5’SS1 and ri12-5’SS2 decreased *STAT2* intron 6 and 12 retention with no significant impact on STAT2 protein levels (**Fig. 3D-H**). By contrast, ASOs ri12-3’SS1 and ri12-3’SS2 increased intron 6 retention under all conditions and increased intron 12 retention with IFN-I (**Fig. 3D,F**), with ASO ri12-3’SS1 having stronger and more consistent effects across conditions. These ASOs efficiently depleted STAT2 protein expression, with ASO ri12-3’SS1 reaching ∼100-fold decrease in protein levels after 24h IFN-I stimulation (**Fig. 3H**). In sum, ASO treatment impacted STAT2 protein expression in a manner broadly consistent with each ASO’s effect on *STAT2* splicing, with ASOs that decrease retention of both introns tending to increase STAT2 protein levels and ASOs that increase retention of one or both introns significantly decreasing STAT2 protein levels (**Fig. 3G,H**). We tested 4 of the strongest ASOs in three additional LUAD cell lines and found largely consistent effects on intron retention and protein expression (**Fig. S2A-C**).

**Figure 3.**
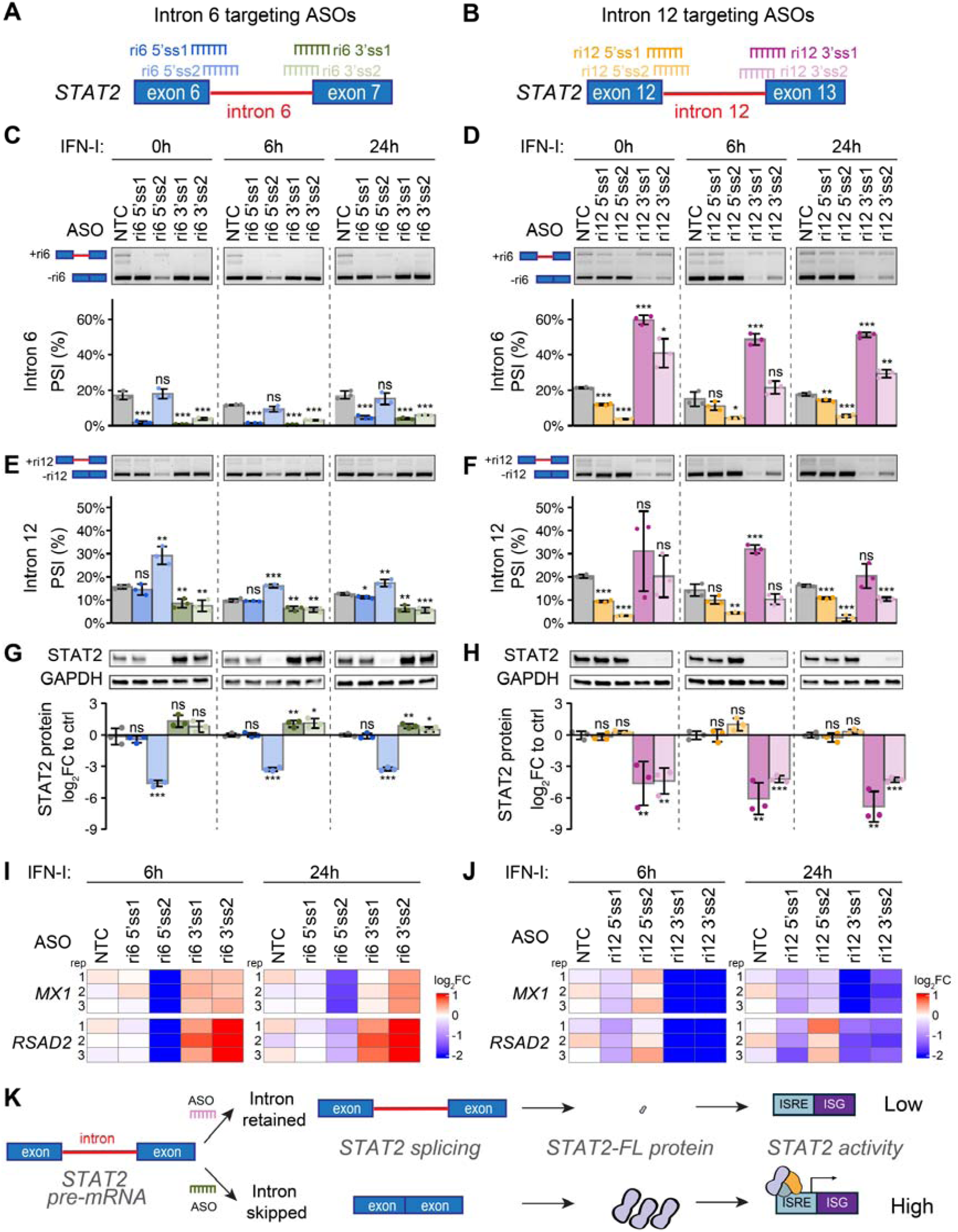
Antisense oligonucleotides targeting *STAT2* retained introns permit dynamic modulation of STAT2 protein expression and IFN-I response. (A,B) Schematic representation of ASOs targeting the splice sites of *STAT2* ri6 (A) and ri12 (B). **(C-F)** Endogenous *STAT2* ri6 (C,D) or ri12 (E,F) retention in A549 cells transfected with ASOs targeting ri6 (C,E) or ri12 (D,F) or non-targeting control (NTC) and treated with 100 U/mL IFN-I for 0, 6, or 24 hours prior to collection is assessed by semi-quantitative RT-PCR with primers that amplify both the skipped and retained-intron isoforms. Representative images along with PSI quantification are shown (*n*=3; mean±sd; student’s *t*-test, \**P*<0.05, \*\**P*<0.01, \*\*\**P*<0.001, ns-not significant, all test relative to NTC-ASO for the same IFN-I timepoint). **(G,H)** STAT2 protein expression in A549 cells transfected with ASOs targeting ri6 (G) or ri12 (H) or non-targeting control (NTC) and treated with 100 U/mL IFN-I for 0, 6, or 24 hours prior to collection is quantified with western blot using an anti-STAT2 antibody and GAPDH as loading control. Representative images along with quantification normalized to NTC-ASO for the same IFN-I time point are shown (*n*=3; mean±sd; student’s *t*-test, \**P*<0.05, \*\**P*<0.01, \*\*\**P*<0.001, ns-not significant). **(I,J)** Expression of interferon-stimulated genes in A549 cells transfected with ASOs targeting ri6 (I) or ri12 (J), or non-targeting control (NTC) and treated with 100 U/mL IFN-I for 6 or 24 hours is quantified by quantitative real-time PCR. Expression is normalized NTC-ASO for the same IFN-I time point (*n*=3; mean±sd; student’s *t*-test, \**P*<0.05, \*\**P*<0.01, \*\*\**P*<0.001, ns-not significant). **(K)** Proposed model for how *STAT2* ri6-and ri12-targeting ASOs impact STAT2 protein expression and IFN-I responses. Increased intron retention depletes full-length STAT2 protein and abrogates the IFN-I response, while decreased intron retention facilitates STAT2 protein expression that permits responses to IFN-I.

Finally, we investigated how ASO treatment affects IFN-I signaling, using the expression of known interferon-stimulated genes (ISGs) as a readout of IFN-I activity. ASOs ri6-5’SS1, ri12-3’SS1, and ri12-3’SS2, which decreased STAT2 protein levels, also decreased expression of the ISGs *MX1* and *RSAD2* after IFN-I stimulation relative to nontargeting control ASO (**Fig. 3I,J**). Conversely, ASOs ri6-3’SS1 and ri6-3’ss2, which increased STAT2 protein levels, increased ISG expression after IFN-I stimulation (**Fig. 3I**). ASOs ri6-5’SS1, ri12-5’SS1, and ri12-5’SS2, which did not impact STAT2 protein expression, did not consistently affect ISG expression after IFN-I stimulation (**Fig. 3I,J**). Altogether, these results demonstrate that *STAT2* splice-switching ASOs can increase or decrease intron retention, which in turn decreases or increases STAT2 protein expression and modulates STAT2 activity in response to IFN-I (**Fig. 3K**).

### STAT2-ri12 can produce a truncated protein which inhibits the type I interferon response

We next investigated whether intron-containing isoforms could produce stable proteins and what their functional consequences would be. Both ri6 and ri12 introduce PTCs that prevent expression of full-length STAT2 protein and could result in truncated STAT2-ri6 and STAT2-ri12 protein isoforms, respectively (**Fig. 4A**). We used AlphaFold3 to predict the structure of these isoforms and found that the STAT2-ri6 coiled-coil domain (CCD), which is disrupted by a PTC, was predicted to extend outwards as a naked alpha-helix, potentially disrupting protein-protein interactions involving the N-terminal domains (**Fig. 4D**). By contrast, the predicted STAT2-ri12 NTD and CCD were structurally similar to those of STAT2-FL (**Fig. 4D**), potentially maintaining canonical STAT2 protein-protein interactions.

**Figure 4.**
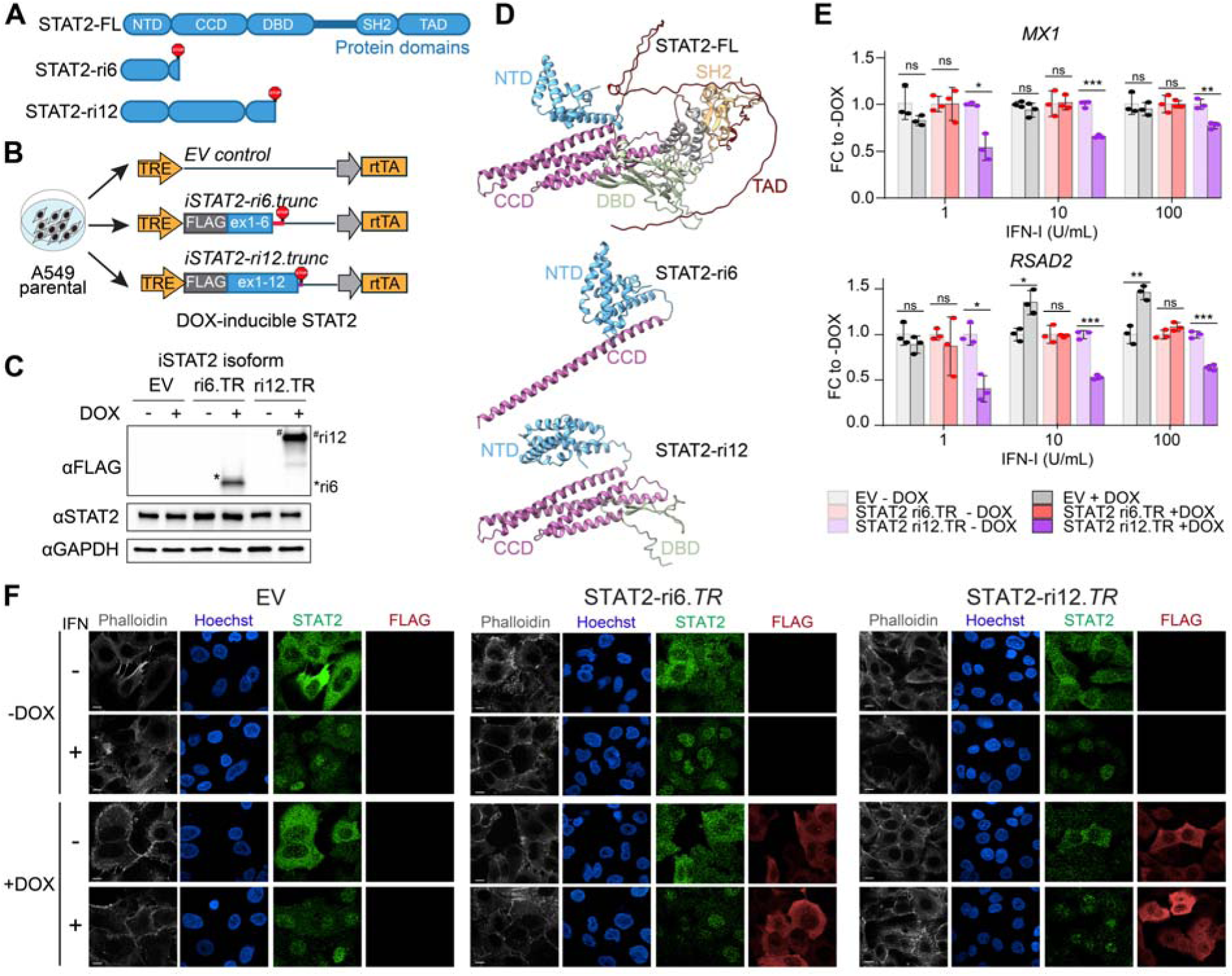
***STAT2* retained intron 6-and 12-containing isoforms encode proteins that fail to respond to IFN-I and differ in their inhibitory capacity. (A)** Domain structures of the full-length STAT2 protein isoform, and the predicted protein isoforms encoded by *STAT2* ri6-containing isoforms or *STAT2* ri12-containing isoforms. NTD = N-terminal domain, CCD = coiled-coil domain, DBD = DNA-binding domain, SH2 = Src homology 2 domain, TAD = transcriptional activation domain. **(B)** Strategy for generating DOX-inducible truncated STAT2 protein isoform-expressing A549 lines. Parental A549 cells expressing wild-type *STAT2* are infected with lentiviral constructs encoding STAT2-ri6.TR or STAT2-ri12.TR. DOX binding activates the reverse tetracycline-controlled transactivator (rtTA) leading to activation of the tetracycline-responsive element (TRE) synthetic promoter that drives construct expression. *iSTAT2-ri6.TR* and *iSTAT2-ri12.TR* are truncated after their respective PTC. **(C)** Expression of FLAG-tagged STAT2 protein isoforms in A549 cells expressing DOX-inducible STAT2-ri6.TR or STAT2-ri12.TR treated with DOX or DMSO for 48 hours is assessed by western blot with anti-FLAG or C-terminal anti-STAT2 antibodies and GAPDH as loading control. Representative images are shown and the expected positions of STAT2-ri6.TR and STAT2-ri12.TR are indicated. **(D)** Protein structures of STAT2-FL, STAT2-ri6, and STAT2-ri12 protein isoforms predicted by AlphaFold 3. Amino acid residues are colored by protein domain from (A). **(E)** Expression of interferon-stimulated genes at 6 hours post-IFN-I stimulation is assessed by quantitative real-time PCR in A549 cells expressing DOX-inducible STAT2-ri6.TR or STAT2-ri12.TR treated with DOX or DMSO for 48 hours, expressed as fold change to the respective DMSO control (mean±sd, *n*=3, student’s *t*-test, \**P*<0.05, \*\**P*<0.01, \*\*\**P*<0.001, ns-not significant). **(F)** Localization of FLAG-tagged inducible STAT2 protein isoforms and STAT2-FL from the endogenous *STAT2* locus is assayed using immunofluorescence microscopy in A549 cells expressing DOX-inducible STAT2-ri6.TR or STAT2-ri12.TR, or EV control, counterstained with phalloidin and Hoechst. Cells were treated with DOX for 48 hours to induce transgene expression or with DMSO as negative control and subsequently were treated with IFN-I (1,000 U/mL) or media-only control for 30 minutes prior to fixation.

We first sought to determine whether *STAT2* truncated isoforms could encode stable proteins. We generated two independent *STAT2*-knockout (*STAT2*-KO) A549 clones using CRISPR/Cas9, and then stably re-expressed cDNAs encoding N-terminally FLAG-tagged full-length STAT2 (STAT2-FL), STAT2 with ri6 (STAT2-ri6), STAT2 with ri12 (STAT2-ri12), or empty vector (EV) control (**Fig. S3A-E**). As expected, no truncated proteins were detected with the C-terminal antibody (**Fig. S3D**); however, despite testing multiple N-terminal STAT2 antibodies, we were unable to find any sufficiently specific for western blotting (data not shown), prompting us to use the N-terminal FLAG antibody for detection. STAT2-ri12 produced a detectable truncated protein (∼50 kDa) detectable with anti-FLAG antibody, but STAT2-ri6 was barely detectable, suggesting that STAT2-ri12 can be expressed at a higher level than STAT2-ri6 (**Fig. S3E**). We stimulated these cell lines with IFN-I and found that only STAT2-FL-expressing cells were able to upregulate *MX1* and *RSAD2* (**Fig. S3F**), demonstrating that neither STAT2-ri6 nor STAT2-ri12 can transduce IFN-I responses.

We next sought to determine how the STAT2 retained intron protein isoforms impact IFN-I signaling in cells expressing both FL and truncated STAT2 isoforms. We generated A549 cells stably expressing doxycycline (DOX)-inducible N-terminally FLAG-tagged STAT2 cDNAs that are truncated after their respective stop codons (referred as *iSTAT2-ri6.TR* and *iSTAT2-ri12.TR*) using a Tet-On system (*64*) (**Fig. 4B,C**). We stimulated cells with IFN-I for 6 hours and compared DOX-induced cells to their respective DMSO controls. Cells expressing STAT2-ri12.TR had reduced ISG expression at all IFN-I concentrations tested compared to control, whereas cells expressing STAT2-ri6.TR showed no difference in ISG expression (**Fig. 4E**). C-terminally FLAG-tagged variants of STAT2-ri6.TR and STAT2-ri12.TR showed similar results (**Fig. S4A-C**).

To examine the possibility that differences in STAT2 protein isoform localization could influence their function, we conducted immunofluorescence microscopy on inducible truncated isoform-expressing cells (**Fig. 4F**). We observed predominantly cytoplasmic signal with a FLAG antibody in iSTAT2-ri6.TR-and iSTAT2-ri12.TR-expressing cells after DOX treatment and no change in localization following IFN-I treatment (**Fig. 4F**). By contrast, STAT2-FL protein detected with a C-terminal STAT2 antibody showed a diffuse and predominantly cytoplasmic signal at baseline, but became intensely nuclear with IFN-I stimulation (**Fig. 4F**), consistent with published reports (*65, 66*). These results suggest that the inhibitory effect of STAT2-ri12 may be mediated by interactions that occur in the cytosol prior to STAT2 nuclear translocation.

Taken together, these data show that isoforms that retain ri6 can encode a truncated protein, STAT2-ri6, which fails to either transduce or inhibit the IFN-I response, whereas isoforms that retain ri12 but not ri6 can encode a truncated protein, STAT2-ri12, which inhibits the IFN-I response.

### *STAT2* retained introns predict response to PD-L1 inhibition

IFN signaling plays a complex role in cancer immunotherapy; IFN signaling is crucial for leukocyte activation, differentiation, and ultimately tumor rejection (*28, 67–70*), but chronic IFN signaling can also lead to immunosuppression and contribute to treatment failure during immunotherapy (*32, 34–36*). To assess how *STAT2* intron retention interacts with the multifaceted role of IFN signaling in cancer, we examined *STAT2* intron retention in LUAD patients from the OAK clinical trial, which compared the PD-L1 inhibitor atezolizumab to docetaxel (*71, 72*) (**Fig. 5A** and **Table S3A**). Briefly, we mapped OAK SR-seq data to our LUAD LR-enriched transcriptome and quantified splicing events using rMATS. We conducted multivariate Cox proportional hazards analysis on atezolizumab-treated LUAD patients with complete metadata to assess the association of overall survival with *STAT2* ri6 and ri12 retention, overall *STAT2* expression, intratumoral PD-L1 IHC, total tumor mutational burden (tTMB), and two IFN signaling scores shown to predict clinical responses to checkpoint blockade: the IFNG.GS score which reflects beneficial IFN signaling and the ISG.RS score which reflects IFN-associated immunosuppression (*35*). *STAT2* ri12 retention predicted improved survival (HR for death: 0.5, 95% confidence interval: 0.30-0.82, *p*=0.006) (**Fig. 5B**). Hazard ratios for continuous variables in Cox proportional hazard models indicate the change in the probability of an outcome with an increment of one unit of the measured variable; here, these results indicate a 50% reduction in the risk of death for every standard deviation increase in *STAT2* ri12 retention pre-treatment. This effect was as strong as a high tTMB (HR for death: 0.4, 95% confidence interval: 0.20-0.82, *p*=0.012). By contrast, PD-L1 staining did not predict overall survival (*p*=0.245), consistent with published OAK data (*72*) (**Fig. 5B**). *STAT2* ri6 retention predicted poorer overall survival (HR for death: 1.67, 95% confidence interval: 1.07-2.61, *p*=0.024). These associations persisted after adjusting for age, sex, and smoking (**Fig. S5A**) and were absent in docetaxel-treated patients (**Fig. 5C, Fig. S5B**), suggesting an immune-specific effect. In the full OAK cohort, ri6 and ri12 retained their prognostic associations (**Fig. S5C,D**).

**Figure 5.**
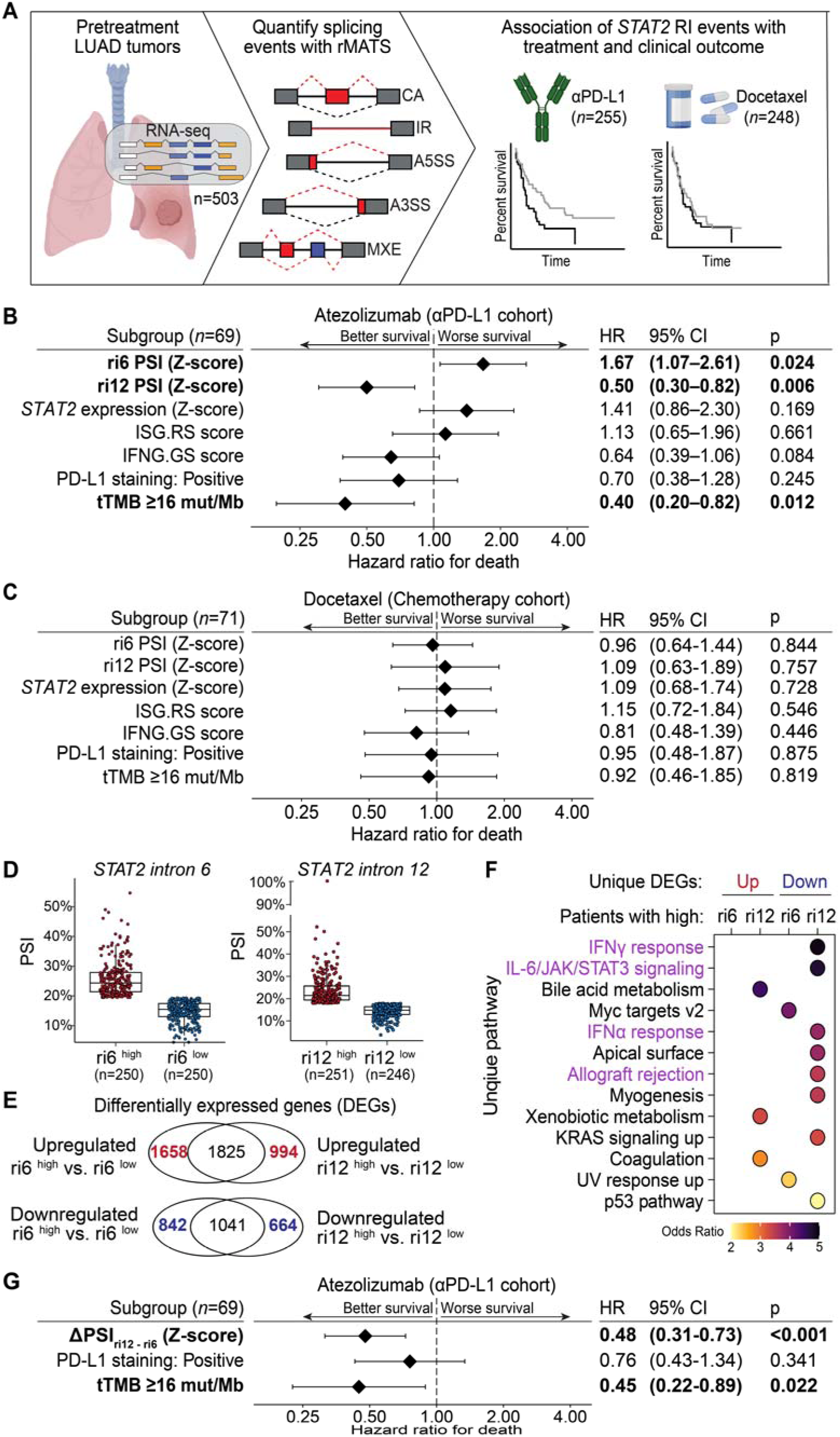
*STAT2* retained introns predict response to atezolizumab. **(A)** SR-seq data from pre-treatment non-squamous non-small cell lung cancer tumors (*n*=503) from the OAK double-blinded Phase III clinical trial was used to quantify alternative splicing events using rMATS, focusing on *STAT2* retained intron events. We then evaluated the link between *STAT2* ri6 and ri12 splicing and survival in patients treated with the PD-L1 inhibitor atezolizumab or chemotherapeutic docetaxel. **(B,C)** Multivariate Cox proportional hazards analysis of overall survival against *STAT2* ri6 and ri12 PSI, *STAT2* gene expression, IFNG.GS (reflecting immunostimulatory/antitumor interferon signaling) and ISG.RS (reflecting interferon-driven immune evasion) scores, PD-L1 expression by IHC stratified as negative (<1% of tumor cells positive) or positive (≥1% of tumor cells positive), and total tumor mutational burden stratified as ≥16 mut/Mb or <16 mut/Mb in the atezolizumab (B) and docetaxel (C) treatment arms. Only patients with all metadata available were included in the model (*n*=69 for atezolizumab arm, *n*=71 for docetaxel arm). All continuous variables were z-scored prior to inclusion in the models and significant covariates are bolded. Hazard ratios and p-values indicate the independent effect of each covariate on overall survival. **(D-F)** OAK patients are classified based on their *STAT2* ri6 or ri12 PSI levels (D) and differential gene expression analysis is conducted between ri6^high^ vs. ri6^low^ patients, and between ri12^high^ vs. ri12^low^ patients (E). Genes upregulated or downregulated uniquely in *STAT2* ri6 ^high^ or ri12^high^ samples are used to conduct pathway enrichment analysis (E,F). Significantly enriched unique pathways from MSigDB (FDR<0.05) enriched among genes uniquely up-or downregulated in *STAT2* ri6^high^ or ri12^high^ samples are shown, with immune-related pathways highlighted in purple (F). **(G)** Multivariate Cox proportional hazards analysis of overall survival against ΔPSI_ri12–ri6_ (calculated as ri12 PSI – ri6 PSI), PD-L1 expression by IHC stratified as negative (<1% of tumor cells positive) or positive (≥1% of tumor cells positive), and total tumor mutational burden stratified as ≥16 mut/Mb or <16 mut/Mb in the atezolizumab treatment arm (*n*=69). ΔPSI_ri12–ri6_ was Z-scored prior to inclusion in the model. Significant covariates are bolded. Hazard ratios and p-values indicate the independent effect of each covariate on overall survival.

Similarly to our cohort and TCGA-LUAD, most OAK patients had similar expression of both retained introns (**Fig. S5E-G**). To identify potential gene expression signatures that could provide mechanistic insight into the opposite predictive effects of *STAT2* ri6 and ri12 on survival, we conducted differential gene expression analysis between LUAD patients in OAK with *STAT2* ri6^high^ (*n*=250) and ri6^low^ (*n*=250), and patients with *STAT2* ri12^high^ (*n*=251) and ri12^low^ (*n*=246), respectively (**Fig. 5D,E**). We identified genes that were uniquely up-or down-regulated in *STAT2* ri6^high^ or ri12^high^ patients, respectively, and conducted pathway enrichment analysis of these unique genes. We filtered for pathways that were specifically enriched among ri6^high^, ri6^low^, ri12^high^, and ri12^low^ groups to identify the distinctive biological consequences of each retained intron. 4 of 5 of the unique pathways most enriched among genes downregulated in patients with *STAT2* ri12^high^ were associated with immune signaling pathways, including IFN-gamma and IFN-alpha signaling (**Fig. 5F**). Repeating these analyses in TCGA-LUAD (*n*=437) showed that genes uniquely downregulated in *STAT2* ri12^high^ patients were also specifically enriched in interferon-gamma signaling (**Fig. S5H**). Taken together, these results suggest that retention of *STAT2* ri12 is associated with suppression of IFN signaling pre-treatment.

The opposing effects of *STAT2* ri6 and ri12 on patient survival raises the intriguing possibility that the difference in inclusion between these introns could be used as a biomarker to predict which patients might derive clinical benefit from atezolizumab treatment. For each patient, we calculated the difference between the inclusion of *STAT2* ri12 and *STAT2* ri6 (ΔPSI_ri12-ri6_) such that higher values reflect more ri12 inclusion relative to ri6 and vice versa. Cox analysis on atezolizumab-treated patients with ΔPSI_ri12-ri6_, PD-L1 staining by IHC, and tumor mutational burden as covariates (*n*=69) showed that ΔPSI_ri12-ri6_ independently predicts improved survival (HR for death: 0.48, 95% confidence interval: 0.31-0.73, *p*<0.001) (**Fig. 5G**). This effect was as strong a predictor of survival as tTMB (HR for death: 0.45, 95% confidence interval: 0.22-0.89, *p*=0.022), which is an FDA-approved pan-cancer biomarker for PD-1/PD-L1 inhibitors. This result suggests that ΔPSI_ri12-ri6_ may serve as a robust immunotherapy biomarker.

## Discussion

Here, we used LR-seq to comprehensively profile alternatively spliced isoforms in LUAD, revealing hundreds of thousands of isoforms, including in genes implicated in interferon responses, and specifically *STAT2* isoforms with retained introns 6 and 12. These retained introns impair expression of full-length STAT2 and may encode truncated proteins with inhibitory function. We showed these introns can predict patient responses to immunotherapy and have developed RNA-based approaches to manipulate their splicing. Beyond these discoveries, our data is a rich resource to be mined by other researchers and highlights the potential of LR-seq to shed light on the complex transcriptomes of human tumors.

Current immunotherapy biomarkers are imperfect (*73, 74*), and there is a pressing need for more and better biomarkers to guide treatment decisions. While splicing signatures have prognostic value in LUAD (*75, 76*), ours is the first study to link specific alternative splicing events—namely *STAT2* ri6 and ri12—to immunotherapy outcomes. The opposing effects of ri6 and ri12 on survival may relate to the ability of STAT2-ri12, but not STAT2-ri6, to encode a dominant-negative protein that inhibits IFN-I signaling. Given that chronic IFN-I impairs immunotherapy efficacy (*34–36*), STAT2-ri12 may suppress this pathway and improve subsequent outcomes. Additional research will be needed to clarify the mechanism and determine whether *STAT2* intron retention can serve as a biomarker for other tumor types or immunotherapeutic modalities.

If *STAT2* intron retention directly influences therapy response, manipulating it may represent a novel therapeutic target. Here, we leverage an FDA-approved ASO chemistry to modulate intron *STAT2* intron retention, which regulates STAT2 protein expression and influences the strength of IFN-I responses. ASOs targeting *STAT2* splicing could enable clinicians to inhibit or promote STAT2 activity in cancer as necessary, providing a more refined degree of control over inflammation than is currently possible. Additional studies to assess the impact of ASO-mediated *STAT2* splicing manipulation alongside immunotherapy *in vivo* are therefore warranted.

While RNA splicing has been studied for decades, the full diversity of spliced isoforms— especially in immune-related genes—has only recently begun to be fully appreciated (*13, 15, 18, 19, 22, 25*). The large number of well-studied genes that have extensive numbers of novel isoforms revealed here by LR-seq highlights the importance of alternative splicing in tumor biology, including antitumor immunity, and may identify additional therapeutic targets. Novel

RNA spliced isoforms and the protein isoforms they potentially encode could carry out distinct and previously unknown functions. Taken together, these results show that LR-seq enables a better understanding of alternative splicing in cancer and may hold the key to breakthroughs that lead to the next wave of therapeutic advances for this disease.

## Acknowledgments

We thank all members of the Anczukow and Palucka labs for helpful discussions and comments on the manuscript and Ankit Patel from ThermoFisher for assistance with ViewRNA probe design. We acknowledge assistance from Microscopy, Genome Editing, and Genome Technologies Sequencing Services at The Jackson Laboratory (JAX) supported in part by NCI grant P30CA034196. We also acknowledge the use of ChatGPT as an assistive editing tool used to condense language during the editing process.

## Author contributions

Conceptualization: RPE, OA, JB, KP; Methodology: RPE, OA, KP, MK, NKL, MB, TW, SP; Investigation: RPE, MB, AS, PY, RB; Visualization: RPE, OA; Funding acquisition: OA, KP, JB, RPE; Project administration: RPE, OA, KP; Supervision: OA, KP, JB; Writing – original draft: RPE, OA, MB; Writing – review & editing: RPE, OA, KP, RB, JB, NKL, MB, SP.

## Data and materials availability

Our splicing analysis pipeline v2.0 has previously been described (*77*) and is freely available on Github (https://github.com/TheJacksonLaboratory/splicing-pipelines-nf). All other data may be found in the paper and associated supplementary materials upon manuscript publication or is available upon request upon manuscript publication. Supplementary Tables will be released upon manuscript acceptance. Raw sequencing files generated in this study will be deposited to the database of Genotypes and Phenotypes (dbGaP) upon manuscript publication.

## Materials and Methods

### Clinical samples

This study was conducted according to the protocol approved by the Institutional Review Board of The Jackson Laboratory for Genomic Medicine (IRB number 2019-RELY-002-HHC). Post-surgical tumor samples from patients with invasive lung adenocarcinoma were collected with appropriate informed consent at Hartford Hospital between the years of 2019 and 2022, and their use was approved by The Jackson Laboratory.

### Human Cell lines

Cell lines were obtained from the American Type Culture Collection (ATCC). A549 lines were cultured in Ham’s F-12 Nutrient Mixture (ThermoFisher Scientific #11765-054) supplemented with 10% FBS (GeminiBio #100-500), 25 mM HEPES (ThermoFisher #15630-080), 2 mM L-glutamine (ThermoFisher #25030-081), and 1% penicillin/streptomycin mixture (ThermoFisher #15140-122). NCI-H1373, NCI-H1573, NCI-H1693, NCI-H1755, and NCI-H1975 lines were cultured in RPMI-1640 medium (ThermoFisher #11875-093) supplemented with 10% FBS (Denville Scientific #C788U23) and 1% diluted penicillin/streptomycin mixture (ThermoFisher #15140-122). All cell lines were grown at 37 °C with 5% CO_2_. For routine subculturing, cells were lifted by incubating cell cultures with 0.05% trypsin-EDTA (ThermoFisher #25300-054) at 37 °C until cells were detached. Cells were routinely tested negative for mycoplasma using the MycoAlert Mycoplasma Detection Kit (Lonza #LT07-318).

### Plasmids

All plasmids were sequence-confirmed using whole plasmid Nanopore sequencing or standard Sanger sequencing (Quintara Biosciences). Primer used for cloning are listed in **Table S4F**.

*pVAX1 transient transfection plasmids*. Full-length STAT2 isoform sequences were derived via fusion PCR from A549 cDNA using Phusion Hot Start II DNA Polymerase (ThermoFisher Scientific #F-549L). All constructs were obtained by sequential fusion PCRs of partial elements of STAT2 obtained from A549 cDNA or gDNA, finally NheI and EcoRI sites were added. All cloning oligos used are listed in Table S4F. The resulting amplicons were digested using NheI-HF (New England Biolabs #R3131) and EcoRI-HF (NEB #R3101) and ligated into the linearized pVAX1 (Addgene #196624) backbone using T4 DNA Ligase (NEB M0202). Empty vector pVAX1 (pVAX1-EV) was generated by restriction digest using NheI-HF (NEB #R3131) and XbaI (NEB #R0145) and ligation of the backbone to itself using T4 DNA Ligase (NEB M0202).

*pLJM1 lentiviral insert plasmids*. Third-generation lentivirus encoding 3xFLAG-tagged *STAT2* isoforms were generated by subcloning each isoform from pVAX1 into pLJM1-Polylinker (Addgene #164143) using T7_F and BGH_R to amplify the insert followed by restriction digest by NheI-HF and EcoRI-HF.

*pINDUCER20 inducible insert plasmids*. All inserts were amplified from pVAX1 using primers AgeI_T7 and PacI_BGH and subcloned into pCR8-ccdb-u1 (gift from Albert Cheng) using AgeI-HF (NEB #R3552) and PacI (NEB #R0547) sites. Inserts were transferred into pINDUCER20 (Addgene# 44012) using Gateway cloning (Invitrogen #11791020).

### Generation of STAT2 knockout cell lines

*STAT2*-targeting gRNA was designed using the CRISPR Design Tool (https://design.synthego.com/<u>#/</u>, Synthego) using the Ensembl GRCh38.p10 human genome build targeting *STAT2* (**Table S4E**). Single guide RNA (sgRNA) with standard chemical modifications (2’-O-methylated nucleotides and phosphothiorate linkages in the first and last three bases) targeting the relevant sequences were synthesized (Synthego). 16 μg of sgRNA was incubated with 20 μg of Alt-R HiFi Cas9 Nuclease V3 (IDT #1081060) for 30 minutes at room temperature. A549 cells were harvested and nucleofected with the sgRNA:Cas9 ribonucleoprotein mixture using the SF Cell Line Solution (Lonza #V4XC-2032) and 4D-Nucleofector X Unit (Lonza) with program CM-130 according to manufacturer instructions. Cells were incubated in complete growth media supplemented with 5 μM Alt-R HDR Enhancer V2 (IDT #10007921). 24 hours after nucleofection, media was removed and replaced with complete growth media without HDR Enhancer. Nucleofected cells were harvested and serially diluted to a concentration of 5 cells/mL and plated into each well of a 96 well plate. Single cell clones were expanded and gDNA was extracted using the QIAamp DNA Mini Kit (Qiagen #51306). The edited region was amplified using primers listed in **Table S4E**, and PCR products were confirmed by Sanger sequencing. Chromatogram files from single cell clones were deconvoluted into constituent alleles with DECODR (https://decodr.org/) using parental A549 cells as control. STAT2 knockout was confirmed via western blot and interferon stimulation followed by RT-qPCR for known STAT2-dependent ISGs *MX1* and *RSAD2* (*78*).

### Generating stable cell line expression STAT2 isoforms

10 μg of lentiviral plasmids were transfected alongside 5 μg psPAX2 (Addgene #12260) and 2.5 μg pMD2.G (Addgene #12259) into HEK293T using Lipofectamine 3000 (Invitrogen #L3000015) according to manufacturer instructions. Lentiviral supernatants were collected at 48 and 72 hours and concentrated as previously described (*79*). Concentrated lentivirus was then resuspended in Opti-MEM (Invitrogen #31985), and stored at-80 °C until further use.

Cells were plated at 4×10^4^ cells/cm^2^and transduced after 2 hours by supplementing media with polybrene (Sigma #TR-1003-G) to a final concentration of 8 μg/mL and adding the desired volume of concentrated lentivirus dropwise. Cells were selected in either 1 ug/mL puromycin (Gibco #A1113803) for lines generated with pLJM1-derived virus or 4 mg/mL G418 (Gibco #10131035) for lines generated with pINDUCER20-derived virus.

### Interferon stimulation

Cells were stimulated with Universal Type I Human Interferon (PBL Assay Science #11200) at indicated doses and durations using complete growth medium as the diluent.

### ASO transfections

1×10^5^ A549 cells were reverse transfected in a 24 well plate with 2′-O-methoxyethyl-phosphorothioated ASO (IDT) at a final concentration of either 100 nM (ri6 5’SS-1, ri6 5’SS-2, ri6 3’SS-1, STAT2 ri6 3’SS-2, ri12 5’SS-1, ri12 5’SS-2) or 50 nM (ri12 3’SS-1, ri12 3’SS-2) using Lipofectamine RNAiMAX (Invitrogen #13778150) according to manufacturer instructions. For all experiments, A549 cells were also reverse transfected with non-targeting control ASO at the same concentration as other ASOs used in the experiment. 72 hours after transfection, cell lysate for RNA or western blot was collected by direct lysis in the plate.

### RNA extraction from tumors and cell lines

High-quality RNA was extracted from samples using the RNEasy Mini Kit (Qiagen #74106). For primary tumor samples, three 0.3 μm tissue sections were cut using a Cryostat from each OCT-embedded tumor and placed directly into 350 μL of RLT Lysis Buffer supplemented with 1% β-mercaptoethanol. For cell lines, cell were lysed by direct addition of 350 μL of RLT Lysis Buffer supplemented with 1% β-mercaptoethanol to the wells directly. Samples were treated with DNase I (Qiagen catalog number) for 1 hour at room temperature following manufacturer instructions. For RNA extracted from primary tumor samples, RNA quality was assessed with the RNA 6000 Pico Kit (Agilent #5067-1513) on a 2100 Bioanalyzer (Agilent). Only samples with an RNA integrity number above 7.0 were selected for sequencing.

### cDNA synthesis and RT-PCR

RNA was extracted as described above and 200-1000 ng of RNA was reverse-transcribed using Superscript III Reverse Transcriptase (Invitrogen #18080093). Semiquantitative RT-PCR was used to amplify 20 ng of cDNA with optimized melting temperatures and cycle numbers using Q5 High-Fidelity Polymerase with High-GC Buffer (NEB # M0491S). Primer sequences are listed in **Table S4A**. PCR products were separated on a 2% agarose gel with SYBR Safe (Invitrogen #S33102) and imaged on a Chemidoc MP Imaging System (Bio-Rad). Band intensities were quantified with ImageLab 6.1.0 (Bio-Rad). The PSI was calculated as the intensity of the included band of the target event divided by the sum of the included and skipped bands of the target event. For each target event, bands of interest were extracted with a Gel Extraction Kit (Qiagen #28704) and Sanger sequenced (Quintara Biosciences) to confirm their identity. Only verified bands were included in the calculation of PSI.

### Quantitative RT-PCR

cDNA synthesized as described above. qPCR was used to amplify 10 ng of cDNA with iTaq Universal SYBR Green Supermix (Bio-Rad #1725120) in 384-well plates (Applied Biosystems #4309849) using a QuantStudio 7 Flex machine (Applied Biosystems). Primers are listed in **Table S4B**. Data was analyzed using QuantStudio Real-Time PCR software. All gene expression was normalized to the housekeeping gene *GAPDH*.

### Western blot analysis

Cells were harvested by direct lysis in the cell culture dish in 2x Laemmli buffer (Bio-Rad #1610737) with 5% (v/v) β-mercaptoethanol pre-heated to 95 °C. Samples were boiled at 95 °C for 20 minutes, with brief vortexing during incubation. Samples were allowed to cool and either frozen at-80 °C or ran on a 4-15% gradient TGX Stain-Free gel (Bio-Rad #4568086). Gels were transferred onto 0.2 um nitrocellulose membranes (Millipore) and blocked in 5% (w/v) milk in Tween 20-TBST (50mM Tris pH 7.5, 150mM NaCl, 0.05% (v/v) Tween 20). Blots were incubated with primary antibody overnight at 4 °C and with secondary antibody for 1 hr at room temperature. A ChemiDoc MP Imaging System (Bio-Rad) was used for infrared or chemiluminescent detection. Blots probed with an HRP-conjugated secondary were incubated in Clarity Western ECL Substrate (Bio-Rad #1705060) for 1 minute prior to image acquisition. Protein expression was quantified using ImageLab 6.1.0 software (Bio-Rad) and normalized first to GAPDH as a loading control and then expressed as fold change to experimental controls. All antibodies used and their respective dilutions are listed in **Table S4G**.

### Immunofluorescence

Cells were cultured in a black 96 well imaging plates (Revvity #6055300) for 48h and fixed by incubating wells with 4% (w/v) paraformaldehyde in PBS. Cells were permeabilized with 0.2% (v/v) Triton X-100 in PBS and blocked with 10% (v/v) goat serum (#a-Aldrich G6767 in 0.2% (v/v) Triton X-100 in PBS. Wells were incubated in primary antibody diluted in blocking solution overnight at 4 °C and detected using Alexa-fluor-488 anti-rabbit or Alexa-fluor-568 anti-mouse conjugated secondary antibodies (Invitrogen) used at 1:500 in blocking solution. Cells were co-stained with Hoechst 33342 (ThermoFisher #H3570, 1:2,000) and phalloidin-AlexaFluor 647 conjugate (ThermoFisher #A22287, 1:1,000) in PBS for 15 minutes at room temperature. Wells imaged on an Opera Phenix High-Content Screening machine (Revvity). All antibodies used and their respective dilutions are listed in **Table S4H**.

### ViewRNA FISH

RNA transcripts were visualized in OCT-embedded lung tumor sections using the ViewRNA Tissue Assay Fluorescence Kits (ThermoFisher #QVT0700). Human STAT2 ViewRNA type 4 probes, human KRT8 ViewRNA type 6 probes, and human STAT2_ri6 or _ri12 ViewRNA type 1 probes were obtained from ThermoFisher. The assay was performed according to the tissue-based ViewRNA assay protocol with formaldehyde fixation and a 20-min protease treatment. ViewRNA probes were detected with Leica TSC SP8 confocal microscope at 40× magnification.

### Long-read RNA-sequencing of tumor samples

Iso-seq libraries were prepared according to Iso-seq Express Template Preparation (Pacific Biosciences) using 300ng of RNA. Full length cDNAs were generated using NEBNext Single Cell/ Low Input cDNA synthesis and Amplification Module (NEB) in combination with Iso-seq Express Oligo Kit (Pacific Biosciences), with 12 cycles of PCR amplification. Amplified cDNAs were purified using ProNex beads. All samples yielded over 160ng of cDNA and were used for library preparation. SMRTbell library preparation including DNA damage repair, end repair and A-tailing and ligation with SMRTbell Adaptors (Pacific Biosciences) were performed following manufacturer recommendations. Loading protocols for each sample were made using PacBio SMRTLink software following manufacturer recommendations. Libraries were sequenced on either a PacBio Sequel II or PacBio Sequel IIe instrument (Pacific Biosciences). Each sample was individually sequenced on a single SMRT cell.

### Short-read RNA-sequencing of tumor samples and cell lines

SR-seq libraries were prepared with a KAPA mRNA Hyperprep Kit (Roche) according to manufacturer’s instruction. First, polyA RNA was isolated from 300ng total RNA using oligo-dT magnetic beads. Purified RNA was then fragmented at 85C for 6 mins, targeting 250-300bp fragments. Fragmented RNA was reverse transcribed with random primers with an incubation of 25C for 10 minutes, 42C for 15 minutes, and an inactivation step at 70C for 15 minutes. This was followed by second strand synthesis and A-tailing at 16C for 30 minutes and 62C for 10 minutes. A-tailed, double stranded cDNA fragments were ligated with Illumina unique adaptors (Illumina). Adaptor-ligated DNA was purified using KAPA Pure beads. This was followed by 10 cycles of PCR amplification. The final library was cleaned up using KAPA Pure Beads (Roche). Quantification of libraries was performed using real-time qPCR (ThermoFisher). Sequencing was performed on an Illumina Novaseq 6000 instrument generating paired end reads of 150bp, targeting 100-200m reads for each sample.

### Long-read RNA data processing

LR-seq files were processed according to the Iso-Seq pipeline provided by Pacific Biosciences at: https://github.com/PacificBiosciences/pbbioconda. Briefly, HiFi CCS BAM files from sequencing runs were trimmed and demultiplexed using *lima* (v2.7.1) with the--ccs and--peak-guess parameters. The resultant reads were quality controlled to remove polyA tails and artifactual concatemers using *isoseq refine* (v4.0.0) with the--require-polya paramter. Reads were assembled into unique isoform clusters using *isoseq cluster2* (v4.0.0) and mapped to the hg38 genome with *pbmm2* (v1.13.0). Finally, mapped isoform clusters were collapsed into non-redundant full-length isoforms using *isoseq collapse* (v4.0.0).

### Long-read isoform filtering and quality control

Isoforms identified by LR-seq were filtered to remove low-quality isoforms and artifacts with the SQANTI3 (v5.2) pipeline (*80*), found at: https://github.com/ConesaLab/SQANTI3. To validate novel splicing junctions with SQANTI3, samples selected for LR-seq were also sequenced with short-read RNA sequencing. Briefly, SR-seq FASTQ files were trimmed using *fastp* (v0.20.0) and aligned to the hg38 genome using *STAR* (*81*) (v2.7.10b) with a modified two-pass method wherein reads were aligned to the hg38 genome in a first pass and the resultant splice junction output files were fed back into a second run with the same samples using the--sjdbFileChrStartEnd option. This method maximizes global sensitivity for detection of novel splice junctions. Long-read isoforms were then classified using the *sqanti3_qc.py* script from the SQANTI3 distribution using the GENCODE v44 comprehensive gene annotation on reference chromosomes (*82*) as the reference transcriptome, CAGE peaks from the refTSS database (*83*) (v3.1), the human polyA motif list from SQANTI3, and splice junction output files obtained from the modified two-pass *STAR* method described above. Isoforms were then filtered manually using the *sqanti3_filter.py* script from the SQANTI3 distribution to remove artifacts and ensure remaining isoforms had sufficient support for novel splice junctions from short reads and transcription start/stop sites from either GENCODE v44 or CAGE peaks from refTSS. For a complete description of filtering parameters used, please refer to the ruleFilter.json file in **Supplementary Materials**. The resultant filtered GTF was annotated using a custom Python script (*annotate_PacBio_GTF.py*) that substitutes Ensembl gene ID’s and human-readable gene names from the reference for the gene ID’s provided by the Iso-Seq pipeline. Additionally, the filtered GTF was annotated and merged with the reference GTF GENCODE v44 using a custom Python script (*collapse_PacBio_Ref_GTFs.py*). Briefly, this script annotates long-read isoforms with Ensembl gene ID’s and human-readable gene names from the reference as above, removes redundant long-read isoforms found in the reference, and removes low-quality reference isoforms that are insufficiently supported. This script was run with the options--utr_5prime 50--utr_3prime 50--transcript_support_levels 1,2,3--annot_levels 1,2. In toto, these options 1) remove long-read identified isoforms that are i) full-splice match to a reference isoform and ii) have transcript start and stop sites within 50 base pairs of the closest reference full-splice match isoform, and 2) remove low-quality reference isoforms that have i) a transcript support level of 4, 5, or NA and ii) an evidence level tag of 3.

### Characterizing long-read isoforms

*Per-sample isoform quantification from LR-seq*. We used *FLAIR* (*84*) (v2.0.0) to quantify isoform counts in LR-seq samples. Briefly, our long-read GTF was converted to BED12 format using the *gtfToGenePred* and *genePredToBed* utilities from the UCSC Genome Browser (*85*) (https://hgdownload.soe.ucsc.edu/admin/exe/) with default options. We converted HiFi CCS FASTQ files from sequencing runs into FASTA format using the command “sed-n’1∼4s/^@/>/p;2∼4p’ “$FASTQ” > “$FASTA”” in bash. We then ran *flair quantify* using the long-read BED12 file using the options--tpm,--stringent, and--check_splice. We queried the raw counts file across each patient sample and considered an isoform detected in a sample if the raw count was ≥1 using a custom Python script.

*Calculating UTR length differences of full-splice match isoforms detected in LUAD*. Isoforms were filtered to include only isoforms categorized as full-splice match or reference match by SQANTI3. The diff_to_TSS and diff_to_TTS parameters from the SQANTI3 classification step were multiplied by-1 to obtain values for the difference in isoform 5’ and 3’ UTR lengths relative to the closest matching reference isoform, such that a positive number indicates a longer UTR in that isoform relative to the closest matching reference isoform and a negative number indicates a shorter UTR in that isoform relative to the closest matching reference isoform.

*Isoform open reading frame prediction and comparison to reference sequences*. We ran the isoform open reading frame prediction tool *CPAT* (*86*) (v3.0.2) on our long-read transcriptome FASTA file using options--top-orf=10 and--min-orf=25. The highest scoring predicted ORF for each isoform was collected and filtered to keep all ORFs with a coding probability score ≥0.364 (the optimal value for human ORFs). These putative ORFs were compared to human reference protein sequences from the UniProtKB/Swiss-Prot database (*40*) using *blastp* (*87*) (v2.14.0+). We calculated ORF identity to the UniProt reference as follows: (2 * matching residues) / (query length + reference length) * 100%. For each ORF, we took the best ORF identity value and rounded with the floor function. Results were grouped by isoform structural category from SQANTI3 and binned as indicated.

*Prediction of isoform coding potential*. The identify isoforms with likely frameshifts/in-frame premature termination codons, we used a custom Python script *_identify_PTC_isoforms.py* using default parameters and human reference protein sequences from the UniProtKB/Swiss-Prot database (*40*) as the reference. This script translates isoforms from genes found in the reference in all three open reading frames and looks for isoforms that have reference protein sequence N-and C-termini that are separated in different reading frames or separated by a stop codon in the same reading frame to identify frameshifts and in-frame premature termination codons, respectively. Isoforms that encode a polypeptide sequence that matches a reference sequence at the N-and C-termini in a single complete ORF are categorized as either exact matches to the reference, or inexact matches with internal mismatches or gaps. Isoforms with N-and/or C-termini missing are flagged as such. For each reading frame defined by a reference N-terminal peptide match, the distance from the stop codon of that reading frame to the last exon-exon junction is calculated. After collecting this data, we used a custom Python script *1_classify_isoforms.py* to output predictions of isoform coding potential. If an ORF from an isoform exactly matched a reference protein sequence, we labeled it as an exact match to the reference; if an isoform’s ORFs inexactly matched one or several reference protein sequences but exactly matched no reference protein sequences, we labeled it as a likely productive isoform with novel internal amino acid sequences. If there were no exact or inexact matches to a reference, we looked for reference ORFs interrupted by frameshifts or PTCs. If all PTC-encoding ORFs had a stop codon more than 55 nucleotides upstream of the last splice junction (*88*), we labeled it as encoding a truncated ORF; if all PTC-encoding ORFs had a stop codon no more than 55 nucleotides upstream of the last splice junction, we labeled the isoform as having a novel C-terminus; if there were a mix of potentially NMD-sensitive and NMD-insensitive ORFs as defined by the stop codon rule, we labeled the isoform as unknown and potentially encoding a truncated isoform or novel C-terminal isoform. If we could not identify matches to any reference ORF sequences at both the N-and C-termini, we looked for matches to reference ORFs at the N-terminus that extended to a stop codon no more than 55 nucleotides upstream of the last splice junction. If such ORFs were found, we labeled the isoform as having a novel C-terminus; otherwise, it was flagged as of unknown coding capacity and a possible fragment.

*Identification of isoforms with retained introns*. To identify isoforms with retained introns, we used a custom Python script *1_identify_intron_retention_events.py*. Briefly, this script collects all exon-exon splice junctions in a GTF, then surveys each isoform to identify exons that wholly contain a splice junction and hence also contain the intervening intron. The script also identifies and removes introns that are retained in the canonical reference isoform from the GENCODE v44 comprehensive gene annotation (*82*).

*Classification of isoforms*. Isoforms were jointly defined according to the classifications from the “*Prediction of isoform coding potential*” and “*Identification of isoforms with retained introns*” algorithms above. Isoforms whose coding classification could not be confidently assigned (coding classification “Unknown”) were removed from further analysis. Isoforms were grouped into the following categories: “Exact match to reference protein” if the coding classification subcategory was “Exact match to reference”, irrespective of intron retention status; “Novel protein sequence” if the coding classification subcategory was “Novel internal sequences” or “Novel C-terminus”, irrespective of intron retention status; “Intron retention leading to protein truncation” if the coding classification was “Truncated” and a retained intron was present; or “Protein truncation without intron retention” if the coding classification was “Truncated” and a retained intron was not present.

### Differential splicing and gene expression analysis

We used a custom in-house pipeline to conduct differential splicing analysis, available at: https://github.com/TheJacksonLaboratory/splicing-pipelines-nf. Briefly, the pipeline was used to trim FASTQ files with *trimmomatic* (*89*) (v0.39) and align them to the human reference genome with the merged long-read/reference GTF using *STAR* (*81*) (v2.7.9a). A percent spliced in (PSI) for each splicing event in each sample was calculated using *rMATS* (*90*) (v4.1.2) utilizing both splice junction read counts and exon/intron body read counts to infer PSI. When applicable, a ΔPSI was calculated wherein ΔPSI = (mean PSI)_case_ - (mean PSI)_control_. Gene and isoform count estimates were made using *Stringtie* (*91*) (v2.1.7) with STAR-aligned BAM files as input and compiled into a counts matrix using a custom Python script. Differential gene analysis was conducted using *DESeq2* (*92*) (v1.44.0) using a Wald test to calculate p-values and the Benjamini-Hochberg method to derive false discovery rate-corrected p-values.

### TCGA splicing analysis and gene expression

BAM files corresponding to genome-mapped RNA-seq files from lung adenocarcinoma patients (TCGA-LUAD) were downloaded from the Genomic Data Commons (accession number phs000178.v11.p8) using the gdc-client executable (https://github.com/NCI-GDC/gdc-client) with a manifest derived by checking options for: Data Type:Aligned Reads, Experimental Strategy:RNA-Seq, Workflow:STAR 2-Pass Transcriptome, and Data Format:bam. BAM files were converted back to raw FASTQ format using *samtools* (*93*) (v1.4.1) to sort BAM files by name (option-n) and the *bam2fastq* utility from *bedtools* (*94*) to convert the sorted BAM file into paired-end FASTQ files. FASTQC (https://www.bioinformatics.babraham.ac.uk/projects/fastqc/, v0.11.9) was used to calculate quality metrics for the resultant FASTQ files. Only samples for which both FASTQ files passed default metrics were considered for further analysis. Splicing analysis and gene expression analysis were conducted using the in-house pipeline described above with a read length of 50 bp and strandedness set to “false” to indicate libraries were unstranded.

### Splicing analysis and gene expression from OAK

FASTQ files from pretreatment tumor samples from lung adenocarcinoma patients and associated metadata were downloaded from the European Genome Archives (accession numbers EGAD00001008549, EGAD00001008550, and EGAD00001007703). Splicing analysis and gene expression analysis were conducted using the in-house pipeline described above with a read length of 51 bp and strandedness set to “first-strand” to indicate libraries were reverse stranded.

### Gene enrichment analysis

We used the EnrichR (*95, 96*) web server at https://maayanlab.cloud/Enrichr/ to conduct all gene ontology, pathway, and cell type enrichment analysis.

### Statistical analysis

For the proportion of novel isoforms stratified by coding sequence impact, significance was assessed with a hypergeometric test taking the *p*-value for over-or under-enrichment as appropriate. For splicing from short-read RNA-seq, significance was assessed using the *rMATS* statistical model (*90*) using the FDR-corrected *p*-value. For comparison of WGCNA-derived gene expression modules with *STAT2* intron retention PSI values, Pearson correlation followed by Benjamini-Hochberg *p*-value adjustment was used. All gene, pathway, and cell type enrichment analysis results were filtered for FDR-adjusted *p*-values<0.05 as reported by EnrichR. For RT-PCR, western blots, and quantitative RT-PCR, data is shown as mean±standard deviation and significance was assessed using an unpaired two-tailed Student’s *t*-test. Survival analysis was conducted using right-censored survival data from OAK and significance was assessed using the Cox proportional hazards model (*97*).

### Graphs and figures

Data was processed using Python or R and plots were generated with R using the ggplot2 library or Microsoft Excel. Figures were made with Abode Illustrator 2023 for Mac incompliance with Nature Publishing Group guidelines concerning image integrity.

## SUPPLEMENTARY FIGURES

**Supplemental Figure 1.**
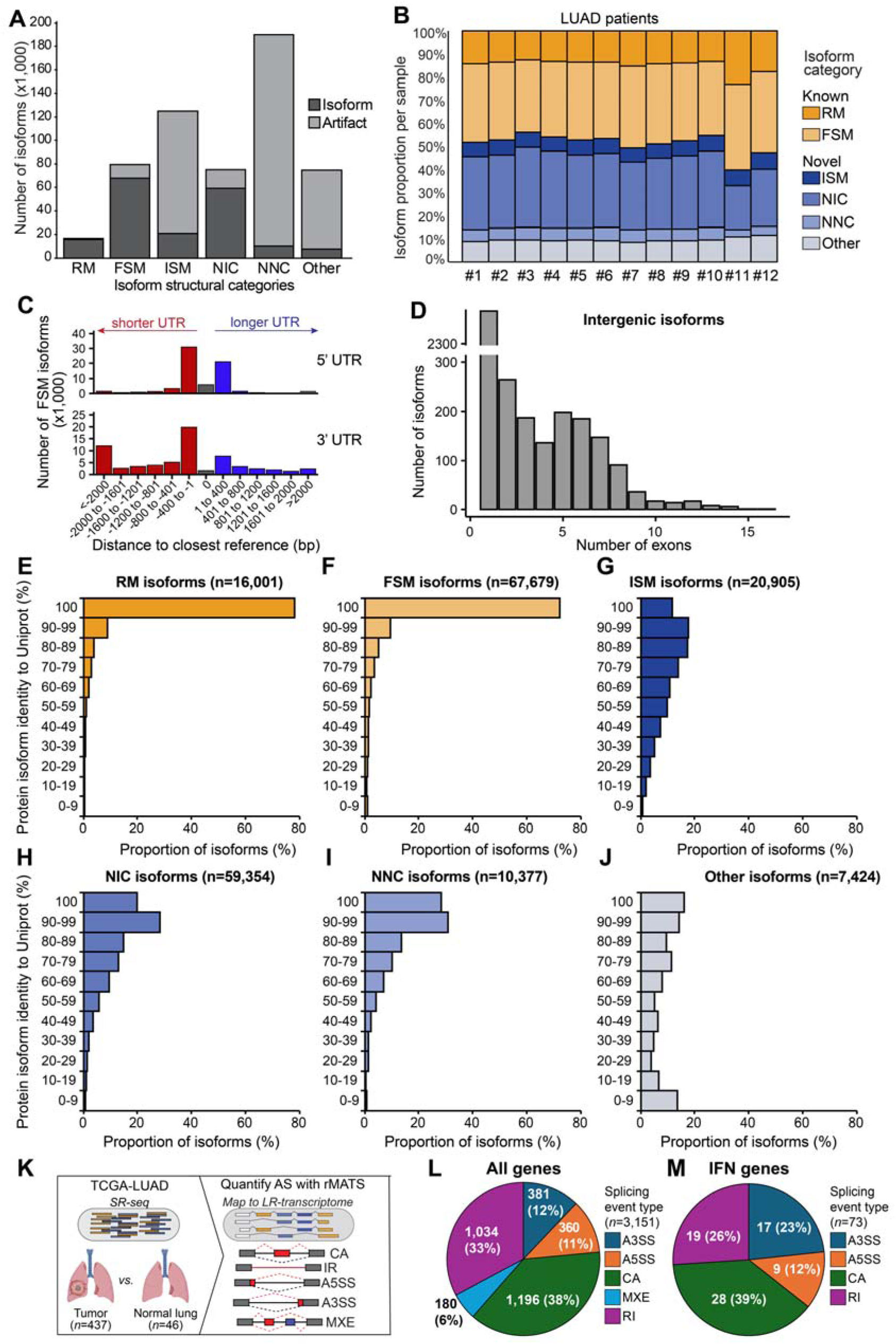
L**R-seq detects novel alternatively spliced isoforms in LUAD.** (related to Figure 1). **(A)** Number of isoforms categorized as isoforms and artifacts by SQANTI3. Isoforms labeled as artifacts were removed. **(B)** LUAD isoforms identified by LR-seq in individual patients shown by structural category. **(C)** Difference in 5’ and 3’ UTR lengths of FSM isoforms identified in LUAD to their closest-matching reference isoforms, binned as shown. A negative value indicates a shorter UTR than the closest reference, while a positive number indicates a longer UTR. FSM isoforms identified in LUAD exhibited shorter 3’ UTRs (median 3’UTR length _iso-ref_ =-84 nt) compared to their closest-matching reference isoforms, with some isoforms losing several kilobases. **(D)** Number of exons in intergenic isoforms. Multiexonic intergenic isoforms may reflect novel spliced genes. **(E-J)** RNA isoforms from each structural category were translated in all three ORFs and compared to reference protein sequences from UniProt (See **Methods** for details) and binned by their identity to the closest match in Uniprot. Protein isoform identity to UniProt is calculated as (2 * matching residues) / (query length + reference length) and expressed as a percentage. **(K)** SR-seq data from LUAD tumors (*n*=437) and normal lung tissue (*n*=46) from TCGA-LUAD is used to quantify alternative splicing events using rMATS. **(L,M)** The proportion of differentially spliced events between tumor and normal samples from TCGA-LUAD from all genes (L) and interferon-related genes (M).

**Supplementary Figure 2.**
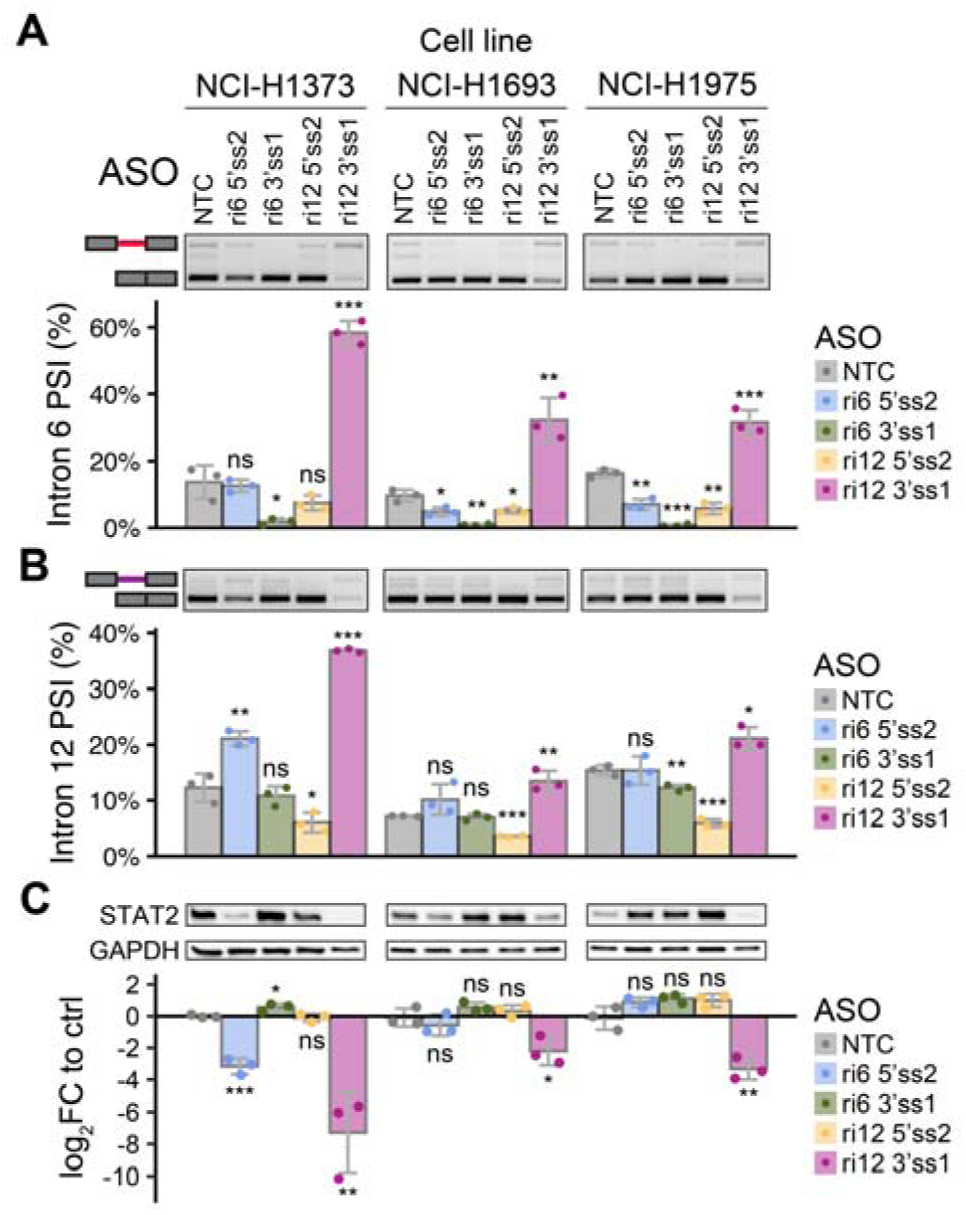
Antisense oligonucleotides targeting *STAT2* retained introns modulate *STAT2* intron retention across cell lines. (related to Figure 3) **(A,B)** Intron retention levels from ri6 (A) and ri12 (B) as assessed by RT-PCR in three LUAD cell lines. All significance tests are with respect to the non-targeting ASO in the respective cell line (mean±sd, *n*=3, student’s *t*-test, \**P*<0.05, \*\**P*<0.01, \*\*\**P*<0.001, ns-not significant). **(C)** STAT2 protein expression, quantified with western blot using C-terminal anti-STAT2 antibody with GAPDH as loading control and normalized as log_2_FC to non-targeting control ASO in three LUAD cell lines. All significance tests are compared to non-targeting ASO in the respective cell line (mean±sd, *n*=3, student’s *t*-test, \**P*<0.05, \*\**P*<0.01, \*\*\**P*<0.001, ns-not significant).

**Supplementary Figure 3.**
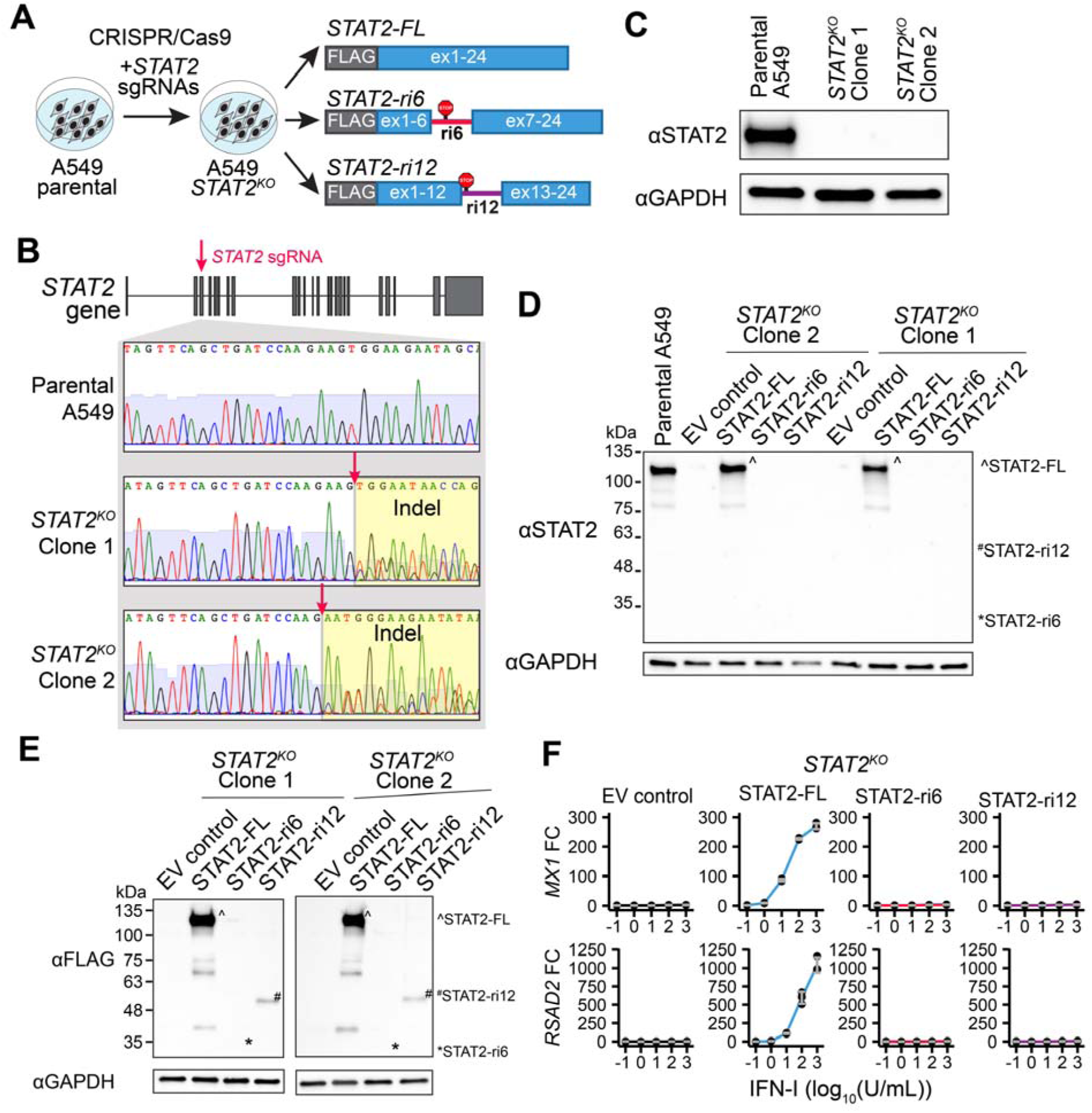
***STAT2-ri6* and *STAT2-ri12* can produce proteins that fail to respond to IFN-I and inhibit full-length STAT2.** (related to Figure 4) **(A)** Strategy for generating stable A549 STAT2^KO^ lines expressing STAT2 isoforms. STAT2^KO^ A549 single cell clones are established using *STAT2*-targeting sgRNA and Cas9, and subsequently infected with lentiviral constructs encoding indicated N-terminally FLAG-tagged *STAT2* isoforms or empty vector control (EV). **(B)** Schematic of the *STAT2* gene showing the location of the *STAT2*-targeting sgRNA used to generate STAT2^KO^ A549 cell lines. Sanger sequencing of parental A549 cells as well as STAT2^KO^ clones 1 and 2 showing the CRISPR/Cas9-derived insertion/deletion (indel) mutations at the target site of *STAT2*-targeting sgRNA. **(C)** Expression of STAT2 protein in parental A549 cells as well as STAT2^KO^ clones 1 and 2 is assessed by western blot using a C-terminal anti-STAT2 antibody and GAPDH as loading control. **(D)** Expression of FLAG-tagged STAT2 protein isoforms in STAT2^KO^ A549 cells stably expressing empty vector control (EV), STAT2-FL, STAT2-ri6, or STAT2-ri12 is assessed by Western Blot with an anti-FLAG antibody with GAPDH as loading control. Representative images are shown, and the expected positions of STAT2-ri6 and STAT2-ri12 are indicated. **(E)** Expression of STAT2 protein isoforms in STAT2^KO^ A549 cells stably expressing empty vector control (EV), STAT2-FL, STAT2-ri6, or STAT2-ri12 is detected by western blot with a C-terminal anti-STAT2 antibody with GAPDH as loading control. Representative images and the expected positions of STAT2-ri6 and STAT2-ri12 are shown. **(F)** Expression of interferon-stimulated genes *MX1* and *RSAD2* at 6 hours post-IFN-I stimulation is assessed by quantitative real-time PCR in STAT2^KO^ A549 cells (clone #1) stably expressing empty vector control (EV), STAT2-FL, STAT2-ri6, or STAT2-ri12, expressed as fold change to *GAPDH*-normalized gene expression of unstimulated controls.

**Supplementary Figure 4.**
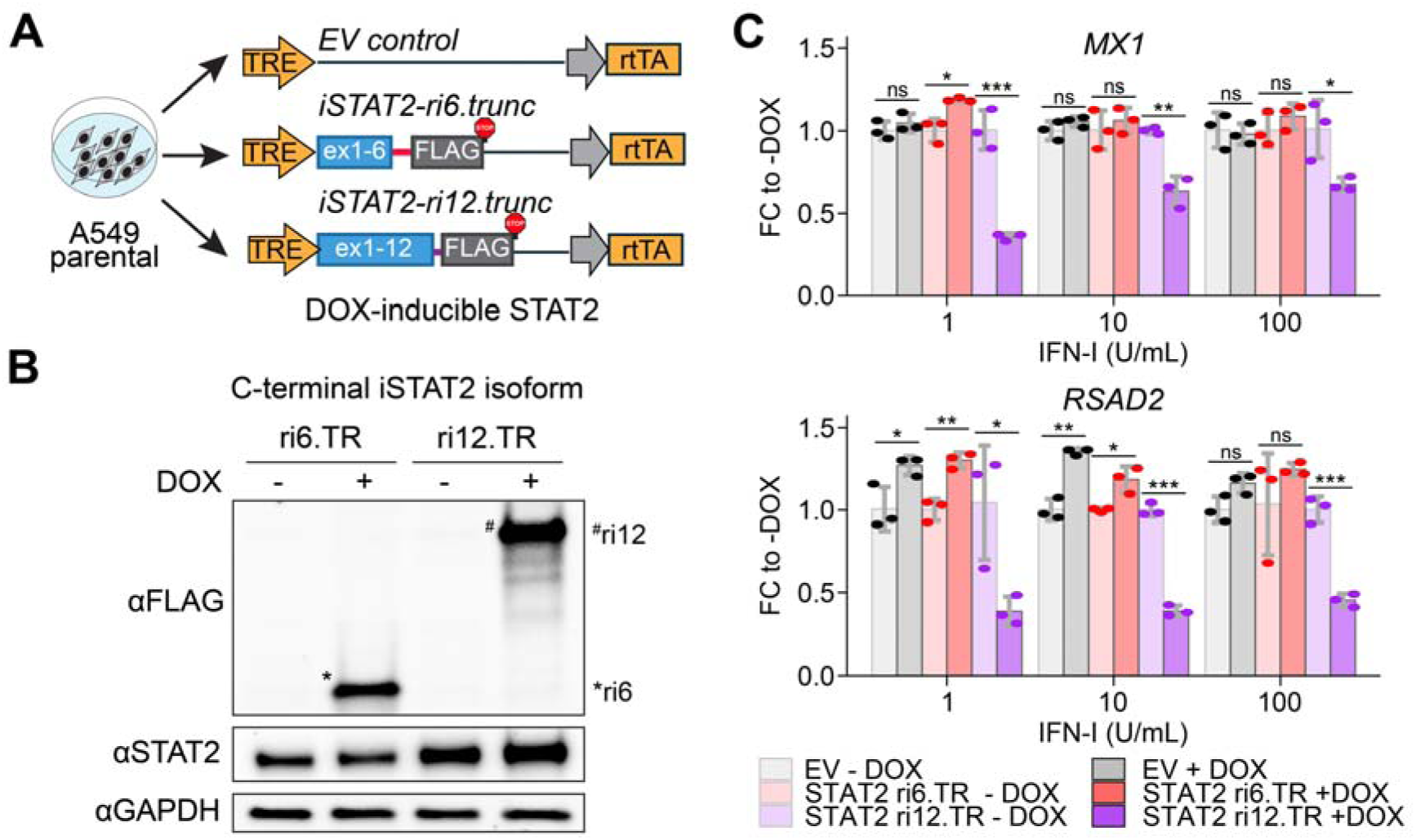
C-terminally FLAG-tagged inducible *STAT2* protein isoforms recapitulate inhibitory capacity of N-terminally FLAG-tagged isoforms. **(A)** Strategy for generating C-terminally FLAG-tagged DOX-inducible truncated STAT2 protein isoforms A549 lines. Parental A549 cells expressing wild-type *STAT2* are infected with lentiviral constructs encoding STAT2-ri6.TR or STAT2-ri12.TR with a 3xFLAG-tag inserted prior to their respective PTC. DOX binding activates the reverse tetracycline-controlled transactivator (rtTA) leading to activation of the tetracycline-responsive element (TRE) synthetic promoter that drives STAT2 isoform expression. **(B)** Expression of C-terminally FLAG-tagged STAT2 protein isoforms in A549 cells expressing DOX-inducible STAT2-ri6.TR or STAT2-ri12.TR treated with DOX or DMSO for 48 hours is assessed by Western Blot with anti-FLAG or C-terminal anti-STAT2 antibodies, with GAPDH as loading control. Representative images are shown and the expected positions of STAT2-ri6.TR and STAT2-ri12.TR are indicated. **(C)** Expression of interferon-stimulated genes at 6 hours post-IFN-I stimulation is assessed by quantitative real-time PCR in A549 cells expressing DOX-inducible C-terminally FLAG-tagged STAT2-ri6.TR or STAT2-ri12.TR treated with DOX or DMSO for 48 hours, expressed as fold change to the respective DMSO control (mean±sd, *n*=3, student’s *t*-test, \**P*<0.05, \*\**P*<0.01, \*\*\**P*<0.001, ns-not significant).

**Supplementary Figure 5.**
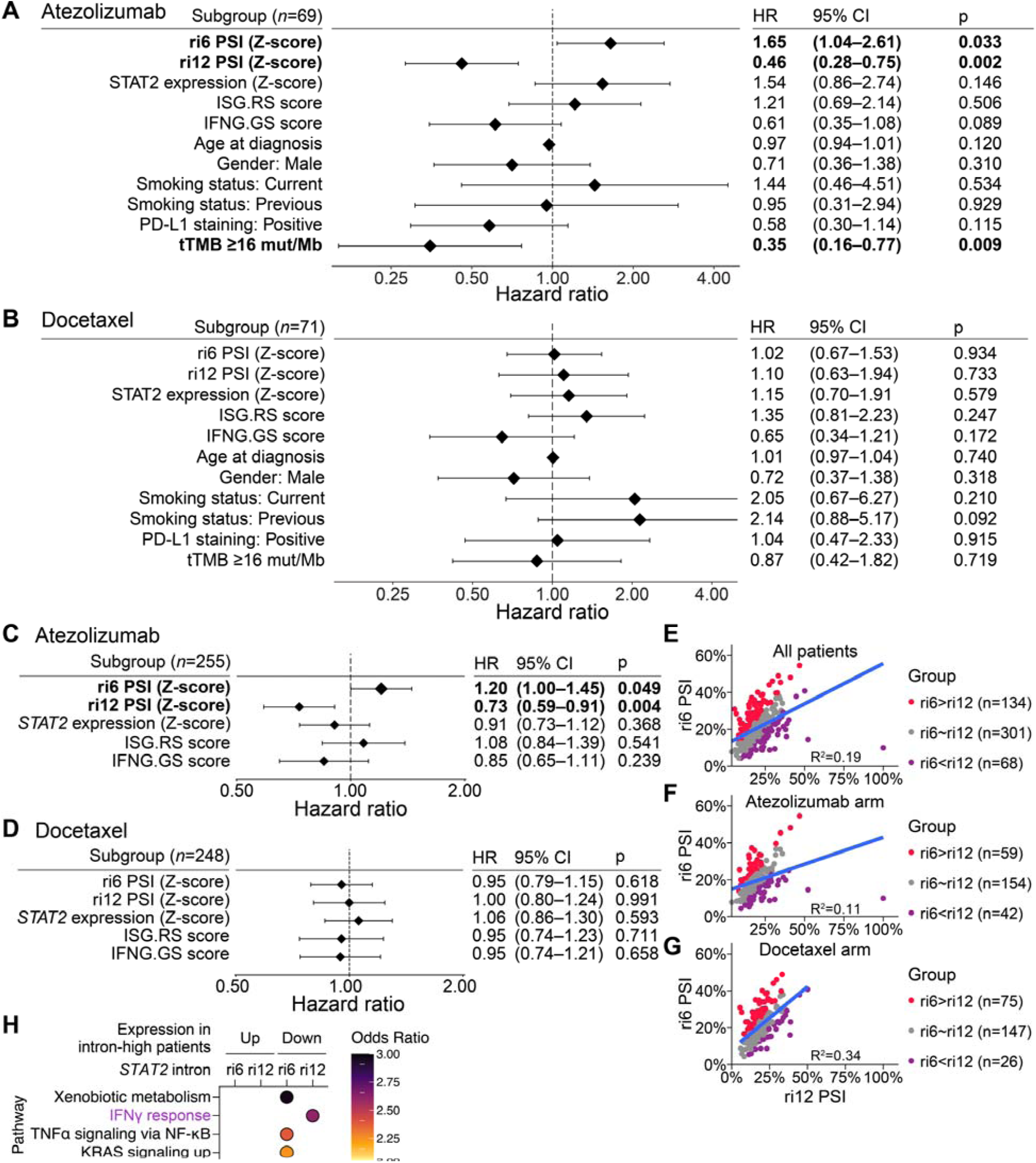
***STAT2* retained introns predict response to PD-L1 inhibition and are associated with unique gene expression signatures across patient cohorts.** (related to Figure 5) **(A,B)** Cox proportional hazards analysis of overall survival among non-squamous patients in the atezolizumab (A) (*n*=69) and docetaxel (B) (*n*=71) treatment arms from OAK against *STAT2* ri6 and ri12 PSI, *STAT2* gene expression, IFNG.GS and ISG.RS scores, PD-L1 expression by IHC stratified as negative (<1% of tumor cells positive) or positive (≥1% of tumor cells positive), total tumor mutational burden (tTMB, stratified as ≥16 mut/Mb or <16 mut/Mb), age, gender, and smoking status (stratified as current, previous, or never). Only patients with all metadata available were included in the model. **(C,D)** Cox proportional hazards analysis of overall survival among non-squamous patients in the atezolizumab (C) (*n*=255) and docetaxel (D) (*n*=248) treatment arms from OAK against *STAT2* ri6 and ri12 PSI, *STAT2* gene expression, and IFNG.GS and ISG.RS scores. **(E-G)** Correlation between retention of ri6 and ri12 in (E) all non-squamous patients from OAK (*n*=503), (F) the atezolizumab arm only (*n*=255), and (G) the docetaxel arm only (*n*=248). Tumors are grouped as follows: ri6>ri12 (ri6 PSI - ri12 PSI ≥5%), ri6∼ri12 (5% > ri6 PSI - ri12 PSI >-5%), or ri6<ri12 (ri6 PSI - ri12 PSI ≤-5%).

